# fuNTRp: Identifying protein positions for variation driven functional tuning

**DOI:** 10.1101/578757

**Authors:** Maximilian Miller, Daniel Vitale, Peter Kahn, Burkhard Rost, Yana Bromberg

**Affiliations:** Department of Biochemistry and Microbiology, Rutgers University, 76 Lipman Dr, New Brunswick, NJ 08901, USA; Columbian College of Arts and Sciences Data Science Program Corcoran Hall, 725 21st Street NW, Washington DC, 20052, USA; Department for Bioinformatics and Computational Biology, Technische Universität München, Boltzmannstr. 3, 85748 Garching/Munich, Germany; Institute for Advanced Study at Technische Universität München (TUM-IAS), Lichtenbergstraße 2a 85748, Garching/Munich, Germany; Department of Genetics, Rutgers University, Human Genetics Institute, Life Sciences Building, 145 Bevier Road, Piscataway, NJ 08854, USA

## Abstract

Evaluating the impact of non-synonymous genetic variants is essential for uncovering disease associations and mechanisms of evolution. Understanding corresponding sequence changes is also fundamental for synthetic protein design and stability assessments. However, the performance gain of variant effect predictors observed in recent years is not in line with the increased complexity of new methods. One likely reason for this might be that most approaches use similar sets of gene/protein features for modeling variant effect, often emphasizing sequence conservation. While high levels of conservation highlight residues essential for protein activity, much of the *in vivo* observable variation is arguably weaker in its impact and, thus, requires evaluation at a higher level of resolution. Here we describe ***fu**nction **N**eutral/**T**oggle/**R**heostat **p**redictor* (***funtrp***), a novel computational method that categorizes protein positions based on the position-specific expected range of mutational impacts: *Neutral* (weak/no effects), *Rheostat* (function-tuning positions), or *Toggle* (on/off switches). We show that position types do not correlate strongly with familiar protein features such as conservation or protein disorder. We also find that position type distribution varies across different protein functions. Finally, we demonstrate that position types reflect experimentally determined functional effects and can thus improve performance of existing variant effect predictors and suggest a way forward for the development of new ones.

## INTRODUCTION

Mapping molecular function or pathogenicity effects of genomic variation is crucial to our understanding of evolutionary, pharmocological, and disease mechanisms. Recent decades have seen significant advances in high-throughput experimentation, as well as growing sophistication in the analyses of the resulting data for research and medical purposes (1–3). However, our understanding of genomic variation is still lacking. For example, separate studies, covering a total of over 7,500 (4,5), have found that less than three percent of known disease-causing variants can actually be deemed pathogenic. On the other hand, known disease-causing variants have been found (6,7) in the (likely) healthy individuals of the 1000 Genome Project (8). Here, a key problem is the absence of an experimental gold standard for identifying disease-causing variants (4). Thus, identifying disease-association of the roughly 10,000 protein sequence changing genetic variants of every individual (9) is like looking for the proverbial needle in a haystack.

Focusing on an arguably better-defined task of finding variants that alter protein function may help; however, variant effects are not all black and white, having a range of outcomes (10). While some variants may only marginally alter ligand affinity, others can induce drastic changes (11). Moreover, while subtle molecular modifications are difficult to detect, they can cause phenotypic changes in concert with other mutation-driven changes (12,13). Experimental techniques like Deep Mutational Scanning (DMS) (14) allow for simultaneous assessment of the effects of hundreds of thousands of variants. DMS combines high throughput sequencing with the ability to create large protein libraries, *i.e.* uniting high throughput selection and high throughput sequencing methods. Still, large-scale mutant library generation is limited by a number of factors, such as bias in sequencing preparation, difficulties in designing accurate and meaningful screening methods (*i.e.* which changes are evaluated), as well as significant time and cost requirements (15,16). Thus, it is infeasible to experimentally assess, for example, the effects of all non-synonymous Single Nucleotide Polymorphisms (nsSNPs) of a given individual, much less a population. However, large-scale mutational fitness landscapes resulting from DMS analyses are an exciting resource for the development of new accurate computational variant effect prediction approaches (17).

Single amino acid substitutions caused by nsSNPs are often associated with specific traits (18–20), diseases (21,22), and pharmacological responses (23). Moreover, targeted mutagenesis of specific protein sites is an essential tool in the synthetic biology toolkit (24). Given the broad range of their possible applications, it is not surprising that many computational algorithms for the prediction of single amino acid substitution effects have been developed (>200; as of January 2018). The different approaches range in algorithm complexity (*e.g.* random forests (25) or meta-servers (26), training/development datasets (*e.g.* cancer (27) or stability changes (28), and gene/protein features used (*e.g.* conservation or protein structure (29–31). However, they still have room for improvement (32–35) and despite their increasing number and complexity, there has, arguably, not been a significant improvement in prediction accuracy over the last decade.

Recently, our collaborators (36) established a new classification of protein (sequence) position types -*Toggle* and *Rheostat* – where mutations in *Toggle* positions were mostly severely disruptive of protein function, while mutations in *Rheostatic* positions had a broad range of effects. We further demonstrated (37) that existing computational predictors fall short of accurately differentiating between neutral and non-neutral mutations in the two position types. Thus, for example at a *Toggle* position, mutations that have been experimentally shown to have no effect on protein function, were often computationally identified as having an effect by most predictors. We further concluded that knowledge of position type could potentially improve prediction accuracy.

In an earlier work, *Toggles* and *Rheostats* were characterized on the basis of the distribution of experimentally validated variant effects per protein sequence position (38). However, experimental evaluation of variant effects is still very limited even in comparison to the number of experimentally annotated protein sequences (*e.g.* 558,590 proteins in UniProtKB/Swiss-Prot, release 2018_09 (39)). Moreover, trivially, for the purposes of computational variant effect predictors, once the variant effect is experimentally determined, its prediction becomes irrelevant. In other words, having to experimentally establish the position type precludes using it as a feature in a variant effect predictor.

Here, we present a new machine learning approach, ***fu**nction **N**eutral/**T**oggle/**R**heostat **p**redictor* (*fuNTRp*), which identifies position types using a curated set of sequence-based features. *funtrp* categorizes protein positions based on the expected range of mutational impacts possible at each position; *i.e.* at *Neutral* positions most variation will have no or weak effect, at *Rheostat* positions – a range of effects is possible, *i.e.* functional tuning, and at *Toggle* positions mostly strong effects are expected. We found that protein regions important for molecular functionality are enriched in *Rheostats* and *Toggles*, with the latter dominating crucial residues (*e.g.* catalytic sites). While these findings are in line with the conservation landscape, we observed lower than expected correlation between conservation and position types, particularly for *Rheostats*. Curiously, we found that the distribution of position types varied across protein classes, slightly differentiating enzymes from non-enzymes and significantly varying between enzyme functional classes. We also showed that position types correlate with experimental effect annotations; *i.e.* we were able to fairly accurately predict mutation effects simply by considering the position type. Combining *funtrp* annotation with outputs of the existing variant effect predictors further improved prediction accuracy.

These findings suggest that knowledge of position types is critical for evaluating functional effects of variants. Thus, *funtrp* predictions could aid the development of improved variant effect prediction methods.

## MATERIAL AND METHODS

The *funtrp* training/development process is detailed in Figure 1. Training datasets are summarized in Supplementary Table S1. All *funtrp* source data/code and software (stand-alone Python and Docker image) are available via DOI and listed in section Availability.

**Figure 1.**
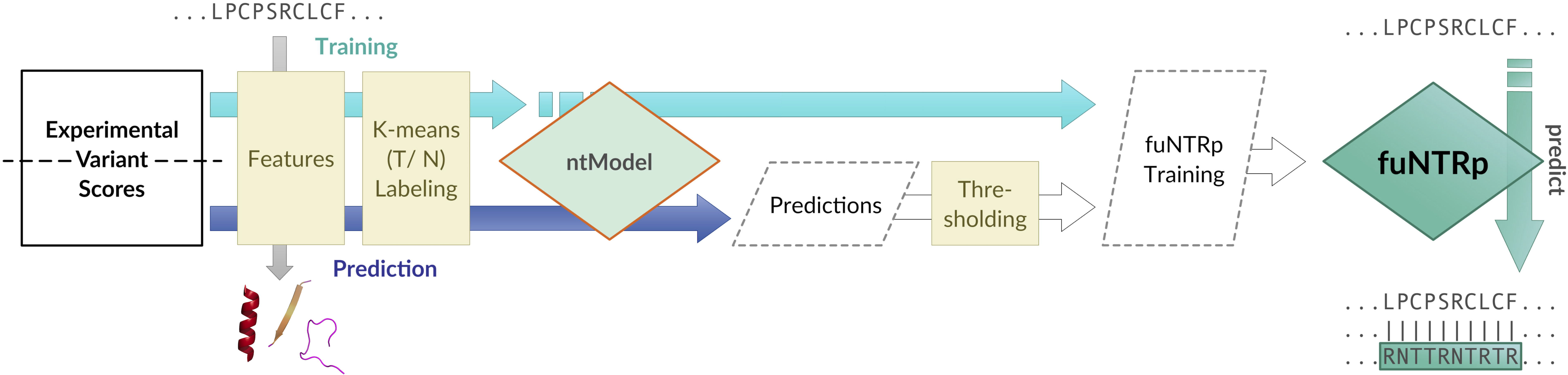
funtrp pipeline. Schematic overview of the *funtrp* pipeline. In training, experimentally measured variant effect scores are extracted for all residues present in selected Deep Mutational Scanning (DMS) datasets. These scores are used in the k-means cluster labeling step to initially label a subset of all (*residue*) positions as either *Toggle* or *Neutral*. Annotated with a computed set of sequence-based features, the subset of cluster-labeled positions is then used to train a Random Forest (RF) (49) classifier (*ntModel*) to predict the not yet labeled positions from the DMS datasets as either *Toggle*, *Neutral*, or *Rheostat*. After filtering, these are combined with the initially (cluster-labeled) positions and used with the same set of sequence-based features to train the final *fuNTRp* model.

### Training datasets and feature extraction

We extracted quantitative deep mutational scanning (DMS) amino acid substitution effect data for five proteins (Table 1) (40–44). The DMS approach generates a large set of mutations and their impact estimates for every evaluated protein-coding gene. For model training, we collected five proteins with the corresponding DMS data available from the literature and meeting a set of stringent requirements: having at least 50 mutated positions, ≥6 variants per position for at least 40% of positions, at least one third of Single Nucleotide Polymorphisms (SNPs) among the variants, wildtype (*wt*) and knockout (*ko*) measurements available, and, notably, available raw datasets in machine readable format (thus omitting data in PDF format and/or not retrievable from corresponding authors; Table 1). To test our model, we collected three additional DMS datasets, covering the PTEN, TPMT, and HSP90 proteins (45,46) which were NOT used in training. Note, that for HSP90 the *ko* effect measures were not directly available; we thus approximated *ko* scores as the mean of the unnormalized effect scores (0.15) reported for variants in eight critical positions, *i.e.* those where any variant resulted in severe functional impact (R32, E33, N37, D40, G94, G118, G121, G123).

For each protein, distributions of effects (scores) of substitutions were standardized to move to zero the *wt* measurements reported in the corresponding publications. All scores (including the *ko* variant scores) were thus transformed to reflect their absolute distance to *wt*, without differentiating beneficial and deleterious mutations (Eqn. 1).

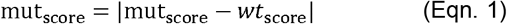

We further computed ten sequence-based features (Table 2 and Supplementary Table S2) for each protein. These features included basic amino acid properties, as well as structural properties generated using a virtualized (Docker) version of PredictProtein (default parameters) (47). Features were chosen based on biological relevance to reflect a broad range of properties associated with protein function.

### Filtering sequence positions

In total, our five proteins comprised 822 amino acids (residues) and 11,130 substitutions with measured effect scores. We removed from consideration the two unknown amino acids (labeled X in sequence), leaving 820 residues. Note that the number of available experimental scores per residue varied between and within datasets. Also note that only half of the available variants (5,423 of 11,130) satisfied the SNP-possible criteria, *i.e.* the observed amino acid substitutions required no more than one nucleotide change with respect to the wildtype amino acid; we did NOT go back to the gene sequence to find the affected codon, but rather designated as SNP-possible any single nucleotide codon to codon changes representing the *wt* and substituting amino acids. As SNPs are more common than multi-nucleotide changes, using only the SNP-possible variants more closely mirrored natural selection patterns. This approach also allowed us to avoid compounding effects of later mutagenesis round mutations, which may have impacted activity more severely.

We removed from any further consideration the 57 positions with fewer than three variant scores as we could not reliably validate any predictions for these positions (7% of 820). With a total of six variants, *Tryptophan* (W) was the amino acid with the least (six) SNP-possible substitutions. Thus, selecting for the first round of training only the positions with at least six SNP-possible variants enabled us to include all *wt* residues, as well as to retain positions with a sufficient number of variants to ensure accurate classification. Thus, we set aside 172 positions (three to five variants; *FewVariants* set) and retained 591 positions (72% of 820) with at least six SNP-possible variants in our dataset, which we labeled the *Clustering* set.

### Toggle and Neutral cluster labeling

We further subdivided the sequence positions in the *Clustering* set into *Neutral* and *Toggle* classes. Note that we previously defined *Toggles* (37) as positions intolerant of any change, while *Neutrals* were new to this work, indicating positions that can tolerate almost all substitutions with no effect on function. Each of the proteins in our set was evaluated separately and only the *Clustering* set positions and variants were considered.

To each protein’s set of experimental variant scores, the protein specific *wt* and *ko* scores were added. This was done to distinguish the effects of variants in the resulting clusters based on the occurrence of these scores. Variants assigned to the same cluster as the *ko* score would be labeled *severe*. Those assigned to the cluster containing the *wt* score would be *no-effect*. All other variants would be labeled *intermediate*. K-means (48) clustering (with K=3) was used to partition each protein variant set into three clusters.

Further, each sequence position *x* was classified (Eqn. 2) into one of two distinct position types (*Toggle* or *Neutral*) on the basis of the distribution of its variant scores among the three clusters. If all but (at most) one variant at *x* were *no-effect*, we labeled *x* as *Neutral* (N; 153 positions). If all but (at most) two variants *severe* we labeled *x* a *Toggle* (T; 66 positions). If none of these two conditions held true, *x* was deemed unknown (372 positions; *Unknown* set).

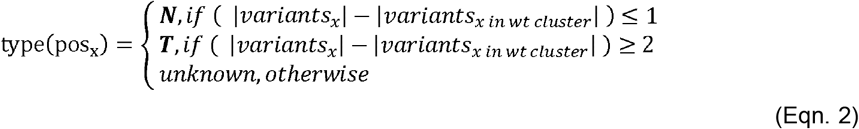

We excluded all unknown positions from *Clustering* and manually refined the remaining positions on the basis of distributions of experimental scores (Supplementary Figure S1). We thus removed six *Toggle* and six *Neutral* positions with noticeably higher score variance and/or different medians of scores as compared to other positions of the same type. We, thus, retained a conservative training set of labeled *Toggle* and *Neutral* positions with comparable variance and medians of experimental variant scores (*ntTraining* set; 207 instances: 60 *Toggles*, 147 *Neutrals*).

### ntModel and Neutral/Toggle scoring

Using the labeled *ntTraining* set we trained a Random Forest (RF) (49) classifier (*ntModel*) to predict *Toggle* vs. *Neutral* position types on the basis of the ten features described above (Table 2). To account for the bias towards the *Neutral* class in the training set, we used over-sampling and trained our model on a balanced input set comprising 414 instances (200% of the unbalanced input). We evaluated the model performance using Leave-One-Out-Cross-Validation (LOO-CV). The model prediction scores were in the [0, 1] range, such that the sum of all type scores equaled 1. The LOO-CV predictions were used to determine prediction score type thresholds, limiting the number of false positive *Toggle* or *Neutral* predictions to ≤3% (Figure 2). Based on this error rate limit, thresholds were set at score ≤0.1 for *Neutral* and score ≥0.8 for *Toggle* predictions.

**Figure 2.**
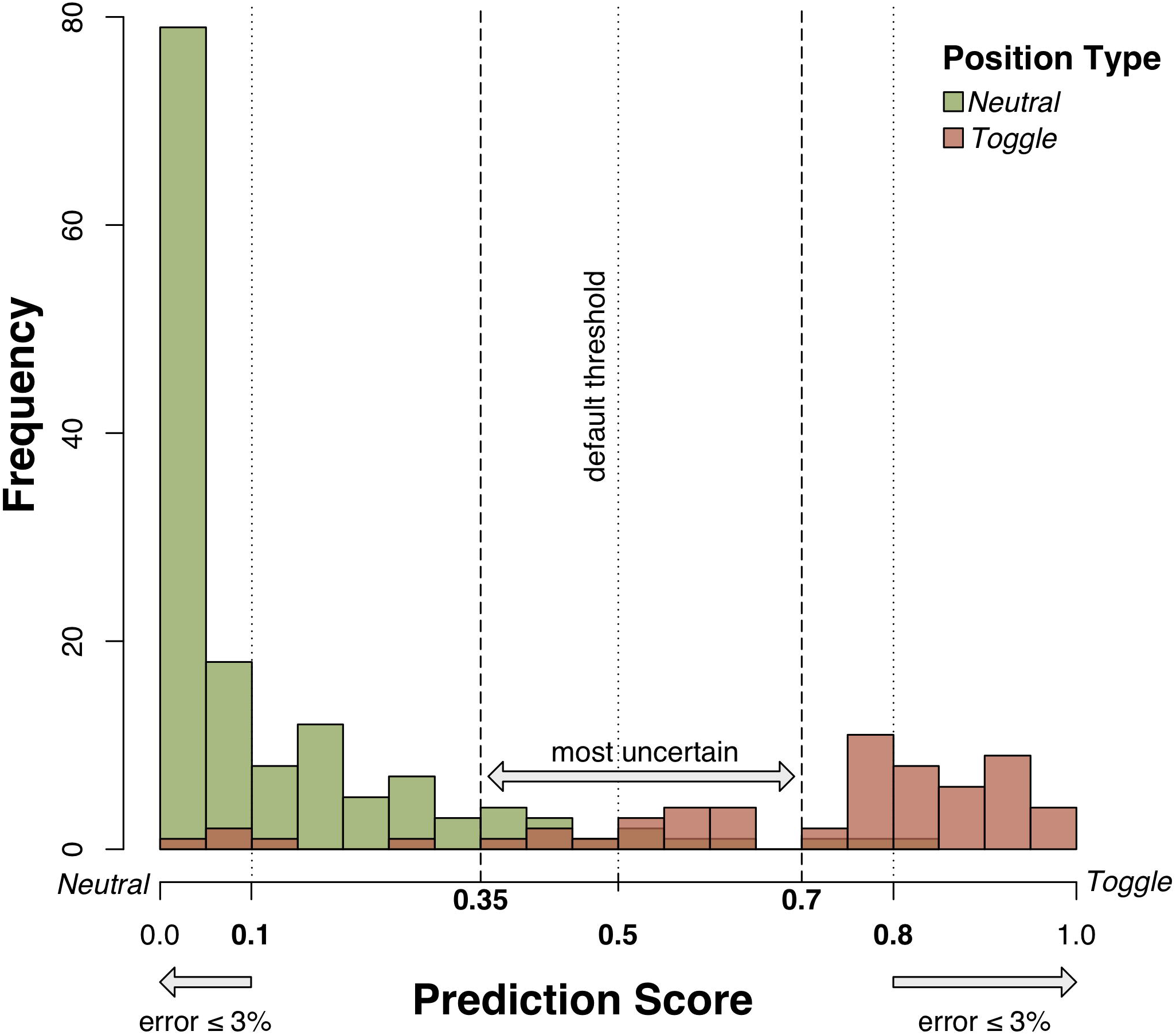
Determination of *ntModel* thresholds. LOO-CV predictions of the *ntModel* were used to determine prediction score type thresholds. Positions with *Neutral* label assigned in the cluster labeling step are shown in green, those labeled as *Toggle* in red. Thresholds were set at score ≤0.1 = *Neutral* and score ≥0.8 = *Toggle*, limiting the number of false positive *Toggle* or *Neutral* predictions to ≤3%. Positions with prediction scores in the range [0.35, 0.7] (containing 50% of all incorrect predictions of the *ntModel*) were defined as *Rheostats*. Remaining positions were excluded from further consideration.

### Defining Rheostats

We assessed the predictions close to the middle (0.5) of our RF classifier prediction range. Here the model exhibited the highest uncertainty in deciding whether the position is a *Neutral* or a *Toggle*. We concluded that positions with prediction scores in that range were *Rheostats* – positions in which mutations can result in a whole range of functionality changes. The *Rheostat* score range was set at [0.35, 0.7] – a range containing 50% of all incorrect predictions of our *ntModel*.

### *funtrp* and residue labeling

The *FewVariants* and the *Unknown* sets comprised 544 (66% of 820) yet-unlabeled positions. We ran the *ntModel* and used score thresholds, as defined above, to assign *N, R, T* predictions to each position. We excluded *ntModel* predictions in ranges (0.1, 0.35) and (0.7, 0.8), leaving only the most reliably predicted positions.

The *ntModel Toggle* and *Neutral* position variant score distributions were compared to those of the cluster-labeled (Step 2, above) positions. We retained only those *ntModel*-*Neutral* positions whose experimental score medians were less than or equal to the highest median score of the clustering-*Neutral* positions. Similarly, the *ntModel-Toggle*s were retained only if their experimental score medians were more than or equal to the lowest median score of clustering-*Toggle*s. We retained only those *Rheostat*s whose medians the were in-between highest clustering-*Neutral* and lowest clustering-*Toggle* median scores. Thus-labeled positions (72 *Neutrals*, 20 *Toggles*, 104 *Rheostats)* were added to the *ntTraining* set to form the *funtrpTraining* set (403 positions: 219 *Neutrals*, 80 *Toggles*, 104 *Rheostats*).

The *funtrpTraining* set was used to train a second RF model, *i.e.* the final *funtrp* model, using the same ten features, over-sampling-based class balancing (806 instances; 200% of the unbalanced input set), and LOO-CV evaluation as in the *ntModel*.

As in the *ntModel*, the per position prediction score for the *funtrpModel* was in the [0,1] range for each position type (N, R, T), such that the sum total of all type scores equaled 1. By default, the position was assigned to the highest scoring type. Performance for both models in this work was reported as accuracy, precision, and recall (Eqn. 3; for each position type, *Y*, at every score cutoff, true positives, TP, are positions correctly predicted as *Y*; false positives, FP, are non-*Y* positions predicted as *Y*; false negatives, FN, are *Y* positions predicted as non-*Y*).

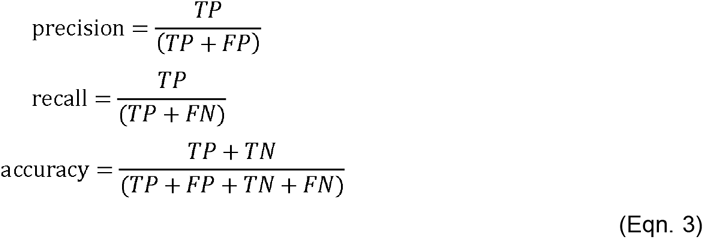

To establish a random baseline for performance evaluation of the final *funtrpModel* we generated random per position predictions. Three scores were randomly sampled from a uniform distribution in range [0,1]. Each score was divided by the sum-total of the three scores, resulting in *Neutral*, *Rheostat* and *Toggle* predictions that add up to 1, analogous to our model. The highest score determined the predicted position type for the random predictor.

### Predicting position types in protein sets

*Neutral*, *Rheostat*, and *Toggle* position types were predicted for various sets of proteins (Supplementary Table S3). All 20,410 manually curated (Swiss-Prot) human proteins were extracted from the UniProt Knowledgebase (UniProtKB release 2018_09) (39). For 5% of these proteins (909; 32 enzymes and 877 non-enzymes) the required set of *funtrp* input features could not be computed. The remaining 19,501 sequences were processed with *funtrp* using *clubber* (50) to distribute computation among multiple High-Performance Cluster (HPC) environments. The subsets of this data were as follows:

1. The *EXPV* set: 1,250 Swiss-Prot enzymes with experimentally validated, unique, unambiguous E.C. (Enzyme Commission) numbers (51).
2. All human enzymes with catalytic site annotations from the M-CSA database (52), which also have binding site annotations in UniProt (94 proteins; 419 catalytic and 214 binding sites).
3. A set of *sahle* spheres (crucial for metal binding; defined as all residues within a 15Å radius sphere from the geometric center of the metal ligand (53)) extracted from 231 transition metal binding protein structures in the PDB (54). PDB structures were mapped to UniProt; *fuNTRp* predictions were available for 230 of these.
4. Disordered (6,309) vs. ordered (13,192) Swiss-Prot proteins (labeled disordered if at least 50% of residues were predicted disordered by the MetaDisorder predictor; MD score threshold of ≥ 0.5 (55)).
5. Proteins containing variants with experimental effect annotations in the Protein Mutant Database (PMD) (56,57) were also collected. We extracted 16,038 variants in 1,224 proteins, along with their SNAP (30), SIFT (29), and PolyPhen-2 (31) predictions of effect from (58). We also extracted precomputed effect predictions of Envision (17), a recently developed method trained on DMS datasets. Note, that Envision predictions where transformed into a binary effect classification by labeling predictions with scores ≥0.9 as *no-effect* and those <0.9 as *effect*. *fuNTRp* was able to make predictions for 1,220 of these proteins, with the remaining four sequences producing error at the feature extraction step. Within this set variants are labeled as either experimentally benign (*neutral*), or having an intermediate (*mild*, *moderate*) or strong effect (*severe*) on protein function. We further extracted from the VarCards database (59) the binary effect predictions (*deleterious* vs. *benign*) of 23 computational predictors. VarCards predictions could be determined for 8,800 PMD variants in 1,042 proteins. For variants with fewer than 23 predictions available, we assigned random prediction scores (uniformly distributed) and generated random binary predictions (>0.5 = *deleterious*, ≤0.5 = *benign*). We defined the per variant *Ensemble Prediction Ratio* as the number of as *deleterious* predictions divided by 23 (the number of predictors). Thus, a ratio above 0.5 (≥12/23) results in an ensemble prediction of *deleterious*, while a ratio below 0.5 (≤11/23) results in a *benign* prediction. Finally, we defined predictions based on the *Ensemble Prediction Ratio* as either correct (the ensemble prediction agreed with the annotated PMD effect) or as incorrect (ensemble prediction in disagreement with PMD effect annotation). Note, that Envision effect predictions were not available for 34 of 8,800 variants (overlap of VarCards and PMD).

### Statistical evaluations

We calculated the standard error of the mean across proteins in different subsets for all three position types. For each subset, we randomly resampled 50% of the proteins (without replacement) in 100 iterations. For each protein we extracted the fraction of each position type. We further computed the standard error across protein subsets of position type fraction means (Eqn. 4; σ = standard deviation of means across subsets; N = total sampling iterations = 100).

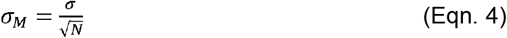

Distributions of feature scores for the three position types were analyzed for similarity using the one-way analysis of variance (ANOVA) test. Prediction performance of variant effect predictors was evaluated per position type using the F_1_ score (Eqn. 5).

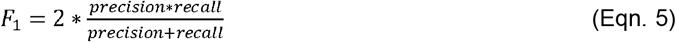

### *funtrp* pipeline implementation

We used a Java based implementation of Random Forest Classification (WEKA, version 3.8) (49,60). We used R (version 3.3.3) (61) for K-Means Clustering, performance and significance evaluations and for visualizations. Protein features were computed using a Docker image of the PredictProtein (47) pipeline (manuscript in preparation). The *funtrp* prediction pipeline requires Python version 3.6 or later and is available as stand-alone version and webservice. References to Docker images applied for this analysis as well as current releases, source code and datasets can be found in section Availability.

## RESULTS AND DISCUSSION

### *funtrp* accurately recognizes position classes

Both RF classifier models were evaluated using LOO-CV (Supplementary Table S4 A,B) at the default cutoff (*i.e.* label of highest prediction score). *ntModel* achieved an overall accuracy of 92.3% (*Neutrals* = 0.94/0.95 and *Toggles* = 0.88/0.85 precision/recall, respectively; Eqn. 3). *funtrp* overall accuracy was 85.1% (*Neutrals* = 0.90/0.91, *Toggles* = 0.88/0.80, *Rheostats* = 0.73/0.77 precision/recall, respectively; Figure 4, solid lines; Eqn. 3). Note that that the higher prediction scores of the *funtrp* model correlated with higher precision, albeit lower recall of the predictions. This performance was significantly above random, which achieved an overall accuracy of 55.5% (*Neutrals* = 0.53/0.32, *Toggles* = 0.20/0.35, *Rheostats* = 0.28/0.36 precision/recall, respectively; Figure 4, dashed lines; Eqn. 3).

**Figure 3.**
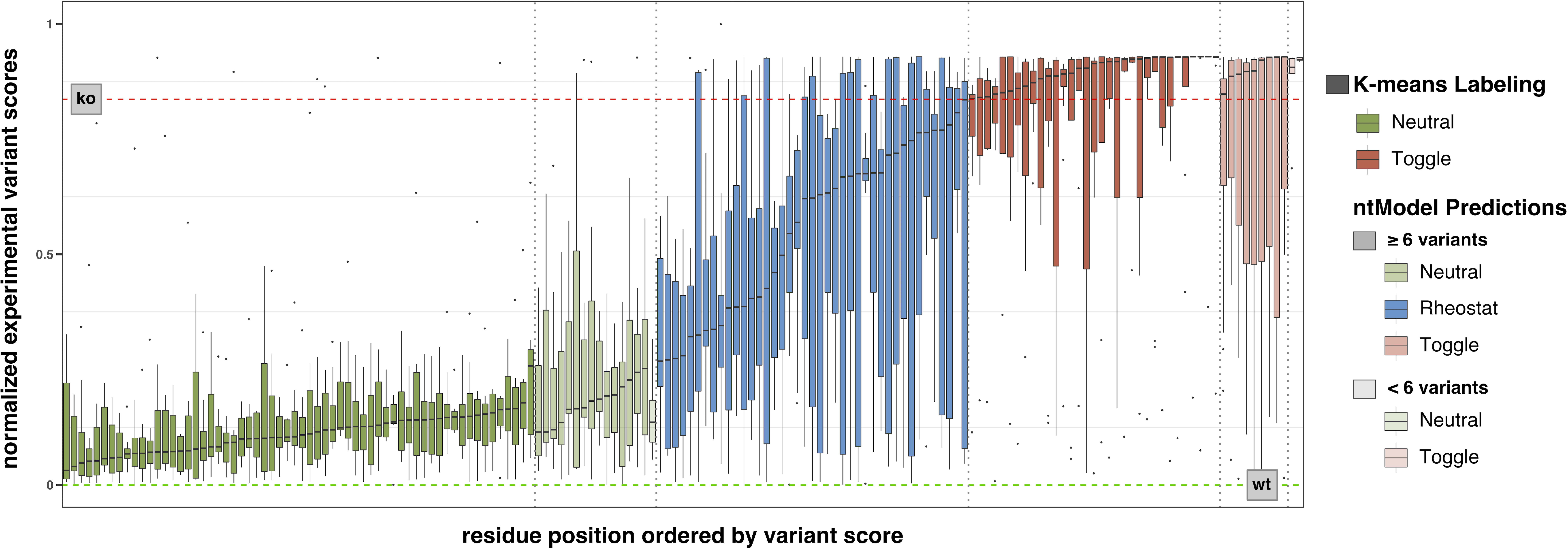
Distributions of experimental effect scores for TEM-1 (E. coli) positions colored by position type. Positions are colored by assigned/predicted position type (green =*Neutral*, blue =*Rheostat*, red =*Toggle*) and ordered by the median of the associated variant score distribution. Positions classified by K-means cluster labeling are shown in opaque colors. Those predicted of the *ntModel* are shown in more translucent coloring based on the number of experimental variants at the respective position. The dashed horizontal lines represent data set specific *ko* (red) and *wt* (green) scores. Positions removed during the manual and automatic refinement steps are not shown. Details for the remaining four proteins are available in Supplementary Figure S1.

**Figure 4.**
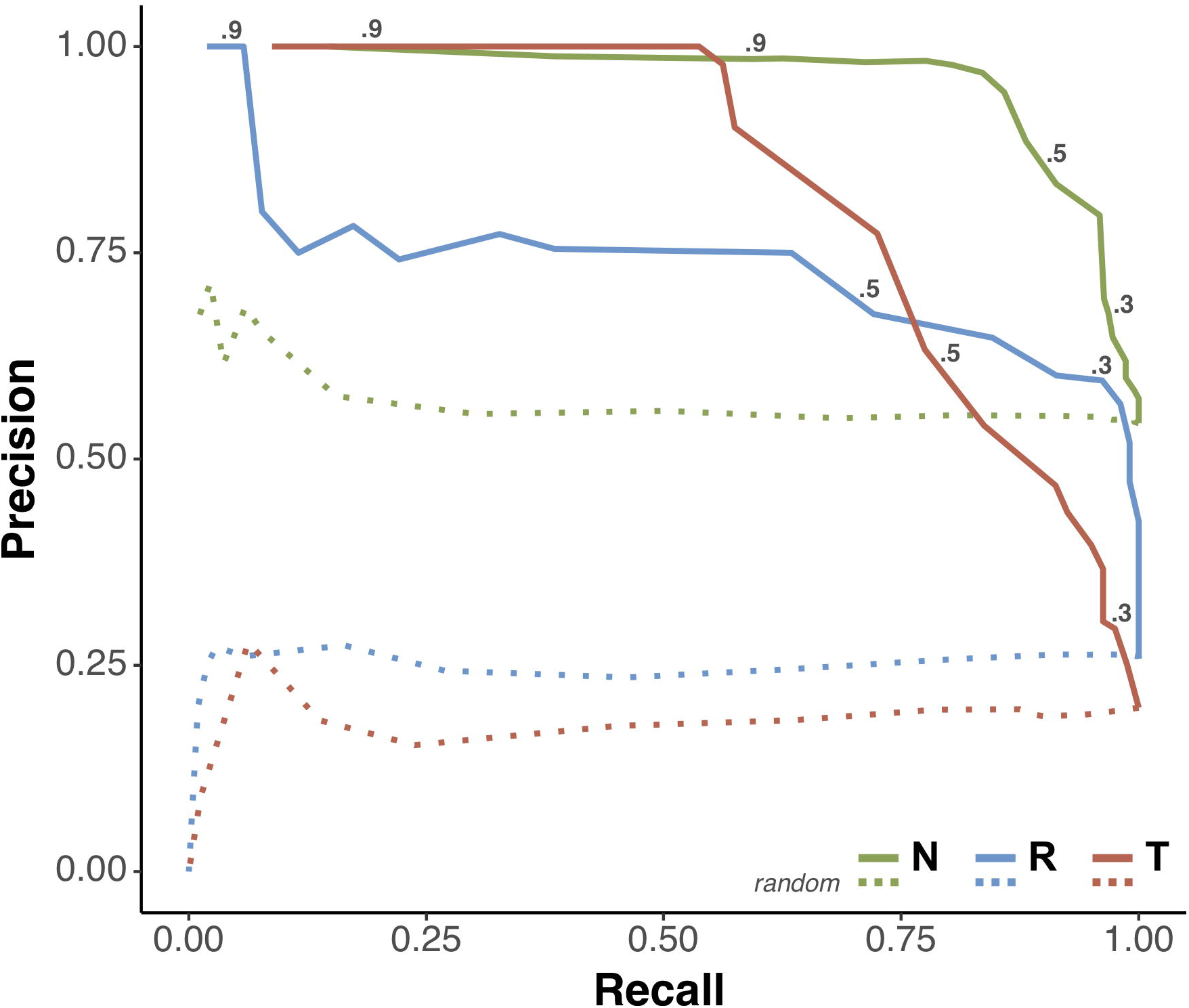
funtrp type classification performance. Precision-Recall curves for LOO-CV predictions of *Neutral* (green), *Rheostat* (blue) and *Toggle* (red) positions for the *funtrpModel* (solid lines) and random predictions (dashed lines). The *funtrpModel* performance for all three position types is indicated for different cutoffs. Performance measures were calculated per position type vs. the other two classes combined for *Neutral*, *Rheostat* and *Toggle* respectively.

The slightly lower performance of the *funtrpModel* (vs. the *ntModel*) in differentiating *Toggles* and *Neutrals* is attributable to the increase set size and less obvious labels of the added positions. A *funtrp* performance discrepancy among classes was expected. After all, while *Toggles* and *Neutral* are explicitly defined types, *Rheostats* are a collection of different position types. As such, they encompass a much larger range/variability in residue properties. For example, in our training set, a position containing three *intermediate* variants would be as much a *Rheostat* if it additionally contained three *no-effect* variants or three *severe* ones.

Additionally, note that truly benign, *no-effect*, mutations are often subjective and always less obvious and more difficult to identify, experimentally or computationally, than *severe* ones. Thus, the differentiation between *Rheostat* and *Neutral* positions is arguably more complex even using experimental data. For *funtrp*, the majority (80%) of the incorrectly predicted *Rheostats* were labeled *Neutral*; more than half of these predictions were also unreliable (scores in the [0.4, 0.49] range). Coincidentally, of the incorrectly predicted *Neutral* positions, 80% were labeled as *Rheostats*.

### Position type labeling is robust across additional proteins

We applied our SNP-possible filtering and clustering - and *ntModel-* based labeling steps, used for generating the *funtrpTraining* dataset, to the three testing DMS datasets, which were not included in model training (Methods). Note that we did NOT apply any manual filtering steps here. We used the labeled positions of these proteins as a *robustness test set*.

We found that the distribution of position types across these sets (PTEN, TPMT and HSP90) (646 positions: 51% *Neutrals*, 27% *Toggles*, 22% *Rheostats*) was very similar to that of the *funtrpTraining* dataset (54% *Neutrals*, 20% *Toggles*, 26% *Rheostats*). The minor discrepancy in *Toggle* and *Rheostat* ratios between the sets was likely due to the fact that no manual filtering was applied to the test set; when manually filtering the training dataset, *Toggles* were affected the most (74% excluded) and *Rheostats* were affected least (31% excluded) (Supplementary Table S1).

We further used *funtrp* to predict the *robustness test set* position types. We found that *funtrp* predictions could re-create the *robustness test set* labels with an average accuracy of 75.9% over all three datasets. The PTEN and TPMT DMS datasets both reporting structural stability, could be re-predicted with accuracies of 79.3% (PTEN; *Neutrals* = 0.84/0.87, *Rheostats* = 0.70/0.78, *Toggles* = 0.87/0.73 precision/recall, respectively) and 84.1% (TPMT; *Neutrals* = 0.91/0.90, *Rheostats* = 0.64/0.71 *Toggles* = 0.88/0.83, precision/recall, respectively). This was comparable to the overall *funtrp* performance of 85.1% (*Neutrals* = 0.90/0.91, *Rheostats* = 0.73/0.77, *Toggles* = 0.88/0.80 precision/recall, respectively). These results on new datasets suggest, that the DMS datasets selected for training were indeed sufficient for our *funtrpModel* training. Larger training sets are often preferred as the number of included feature-observation combinations is thus increased, resulting in more complete models. However, adding more data can also introduce new biases, increase noise, and potential errors; it is thus often accompanied by slight changes in model performance. Thus, as expected, the results of re-prediction of our position type labels for both PTEN and TPMT DMS datasets suggest that adding either protein to training would not drastically change our model performance. However, the HSP90 label reassignment could only be done with 70% accuracy (*Neutrals* = 0.90/0.73, *Rheostats* =0.34/0.62, *Toggles* = 0.73/0.67 precision/recall, respectively). As opposed to directly measuring the effect of mutations on protein function and/or structure, the HSP90 dataset reflects how mutants in Hsp90, a chaperone protein, impact the growth rate of budding yeast. This reduced performance is thus very likely due to two critical experimental design decisions. First, chaperones assist in correct (un)folding of other proteins. Thus, the measured and reported effect is not directly coupled to the mutagenized protein function, but rather to its ability to facilitate function of other proteins. Second, the reported fitness measure of overall growth rate is more removed than what was evaluated for other proteins in our set, *i.e.* ligand binding affinity or protein stability. These observations can likely explain why the re-labeling of HSP90 could lead to the observed model variability and highlight the need for standardized selection of experimental training data.

### Individual sequence-based features are not sufficient to describe position types

Using the ReliefF (62) feature selection algorithm we ranked the importance of *funtrp* features for labeling sequence positions in Swiss-Prot (Table 2). As expected, evolutionary conservation was ranked most important. However, the assigned weight was only slightly higher than other important features: protein disorder, solvent accessibility, and residue flexibility. These results suggest that none of these features alone could explain the predicted position types.

Conservation is widely used as an approximation for residue importance (63,64); *i.e.* the more conserved a residue is, the higher the likelihood that its substitution by another amino acid will result in a function disruption. We compared conservation scores (defined by ConSurf (65) for all positions of experimentally verified enzymes (EXPV). As expected, these were significantly different between the three position types (Figure 5; medians in bold; ANOVA p-value < 2e^−16^). ConSurf scores are normalized by default, so that the average score over all residues of one protein is zero, and the standard deviation is one; here, lower scores indicate more conserved residues. *Toggle* positions were predominantly conserved while *Neutral* positions were for mostly non-conserved. *Rheostats*, however, were in-between the other position types and often showed similarly high conservation as the *Toggles*.

**Figure 5.**
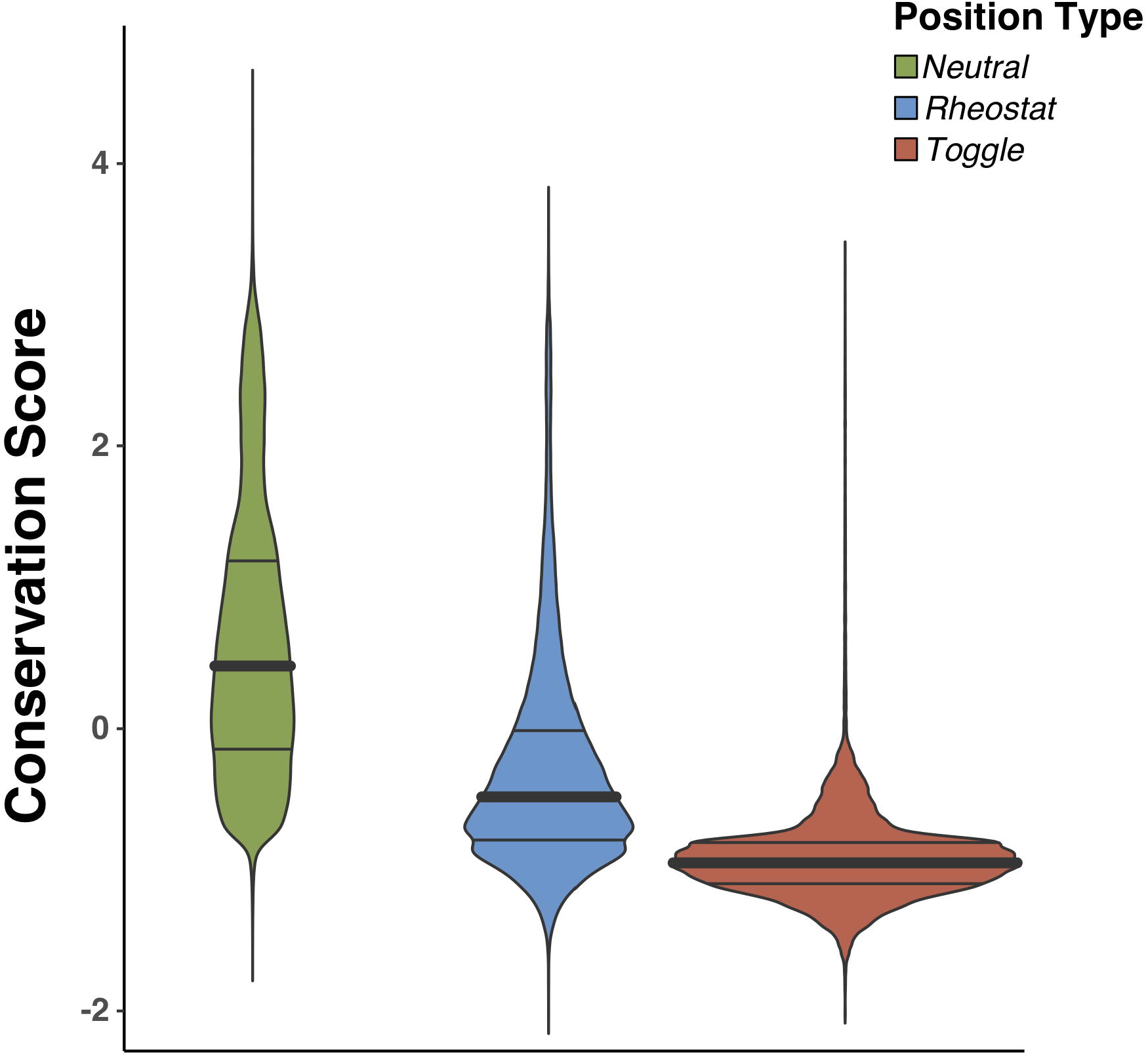
Conservation of position types. Density distributions of evolutionary conservation (ConSurf) compared between position types. ConSurf predictions scores are by default normalized such as 0 depicts the average score over the entire protein and standard deviation is |1|). Distribution medians are highlighted in bold.

To further establish how well a predictor for position types could perform using conservation alone, we computed the number of positions in Swiss-Prot proteins that could be correctly identified as a *funtrp Rheostat, Toggle*, or *Neutral* at a fixed cutoff. The lowest cutoff for *Neutrals* was selected by taking the mean of the distribution medians of *Neutral* and *Rheostat* conservation scores. Similarly, the highest cutoff for *Toggles* was at the mean of *Rheostat* and *Toggle* conservation score medians. *Rheostats* were assigned all other conservation scores. The overall accuracy for this thresholding was 61% (*Neutrals* = 0.80/0.70, *Toggles* = 0.45/0.80, *Rheostats* = 0.44/0.39 precision/recall, respectively; Supplementary Table S5);

Thus, evolutionary conservation-despite being the highest-ranking feature - was not representative of position types. Furthermore, none of the remaining features was likely to perform better than conservation indicated by their consistently lower ReliefF rankings (Table 2). Moreover, arguably, for a given position in a given protein establishing the conservation thresholds for each of the three classes would be infeasible. Note, that we observed the same trends for the training dataset (*funtrpTraining*) for *funtrp* (Supplementary Figure S2).

### Position type profiles differ across protein classes

Swiss-Prot (Figure 6A) enzymes had proportionately more *Toggle* and fewer *Neutral* positions than non-enzymes. However, there was no difference in the number of *Rheostats* between enzymes and non-enzymes. As *Rheostats* allow for functional and evolutionary flexibility while adapting to different environments, the latter result is expected. The increase of *Toggles* in enzymatic proteins, *i.e.* positions critical for defining protein activities: active sites, ligand specificity, *etc.*, is very likely due to enzymes having evolved to implement a set of very specific functionalities. These very mutation sensitive key positions are thus enriched in comparison with non-enzymatic proteins. Note that this increase in *Toggles* at the expense of the reduction in *Neutral* sites is unlikely due to resolution limits of the *funtrp* predictor, as this would likely produce fewer *Rheostats* to account for an increase in *Toggles*.

**Figure 6.**
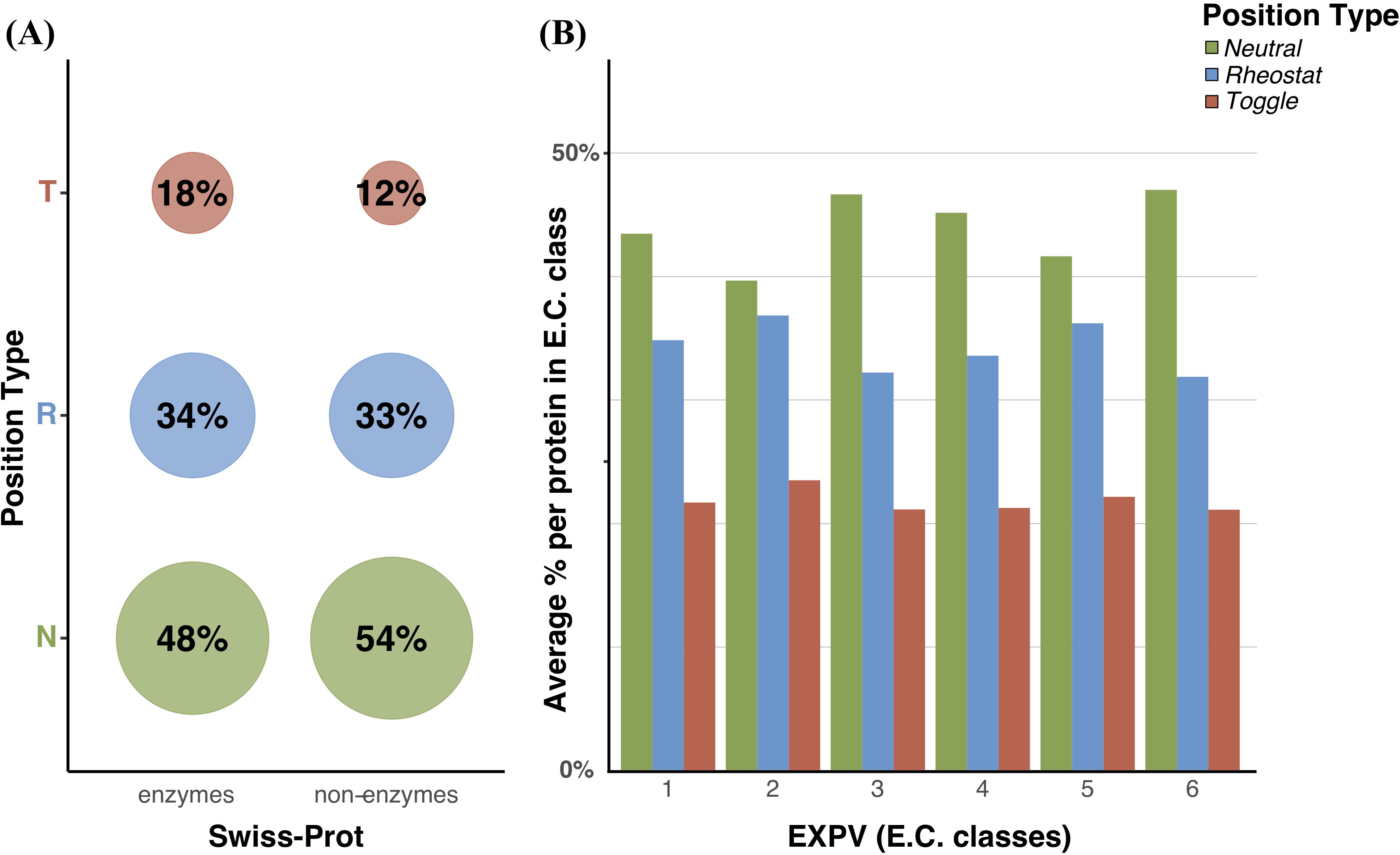
Distribution of position types per protein class. Distributions are based on entire Swiss-Prot (**A**) and EXPV sets (**B**). Colors are according to position type (green =*Neutral*, blue =*Rheostat*, red =*Toggle*). Percentages in (**A**) are rounded off and thus do not add up to 100%. Fractions in **(B)** are averaged on a per-protein basis. Mean fractions of position types in **(B)** differ significantly among enzyme classes based on the standard error of the mean: 1 (N= 6.0E-04, R=6.4E-04, T=5.8E-04), 2 (N=3.7E-04, R=4.1E-04, T=2.4E-04), 3 (N=5.2E-04, R=4.2E-04, T=3.6E-04), 4 (N=9.2E-04, R=1.0E-03, T=7.6E-04), 5 (N=1.5E-03, R=1.1E-03, T=1.2E-03), 6 (N=8.9E-04, R=9.5E-04, T=9.1E-04).

We further compared the mean per-protein fraction of position types between the six main enzyme classes (with corresponding E.C.s) of the experimentally annotated EXPV set: Oxidoreductases (1), Transferases (2), Hydrolases (3), Lyases (4), Isomerases (5) and Ligases (6). Although the general trend of *Neutrals>Rheostats>Toggles* was maintained across all enzyme classes, the classes differed significantly in the actual fractions per position type (Figure 6B). For all EC classes, *Toggle*s made up less than a quarter of all positions per protein, suggesting that enzymes are robust to mutation. We observed similar trends for the full Swiss-Prot dataset (Supplementary Figure S3), with slight differences in position type distributions likely explained by the latter dataset sequence redundancy. Note that the EXPV proteins are experimentally annotated and, thus, tend to be less redundant (98% of the sequences are <90% sequence similar).

### Distribution of position types varies by residue function

We compared the distribution of position types for catalytic sites, binding sites, and *other residues* in Swiss-Prot enzymes (Figure 7A). Note, that here we included only the 47 proteins containing both binding and catalytic sites, which were non-overlapping, *i.e.* annotated in different positions of the protein.

**Figure 7.**
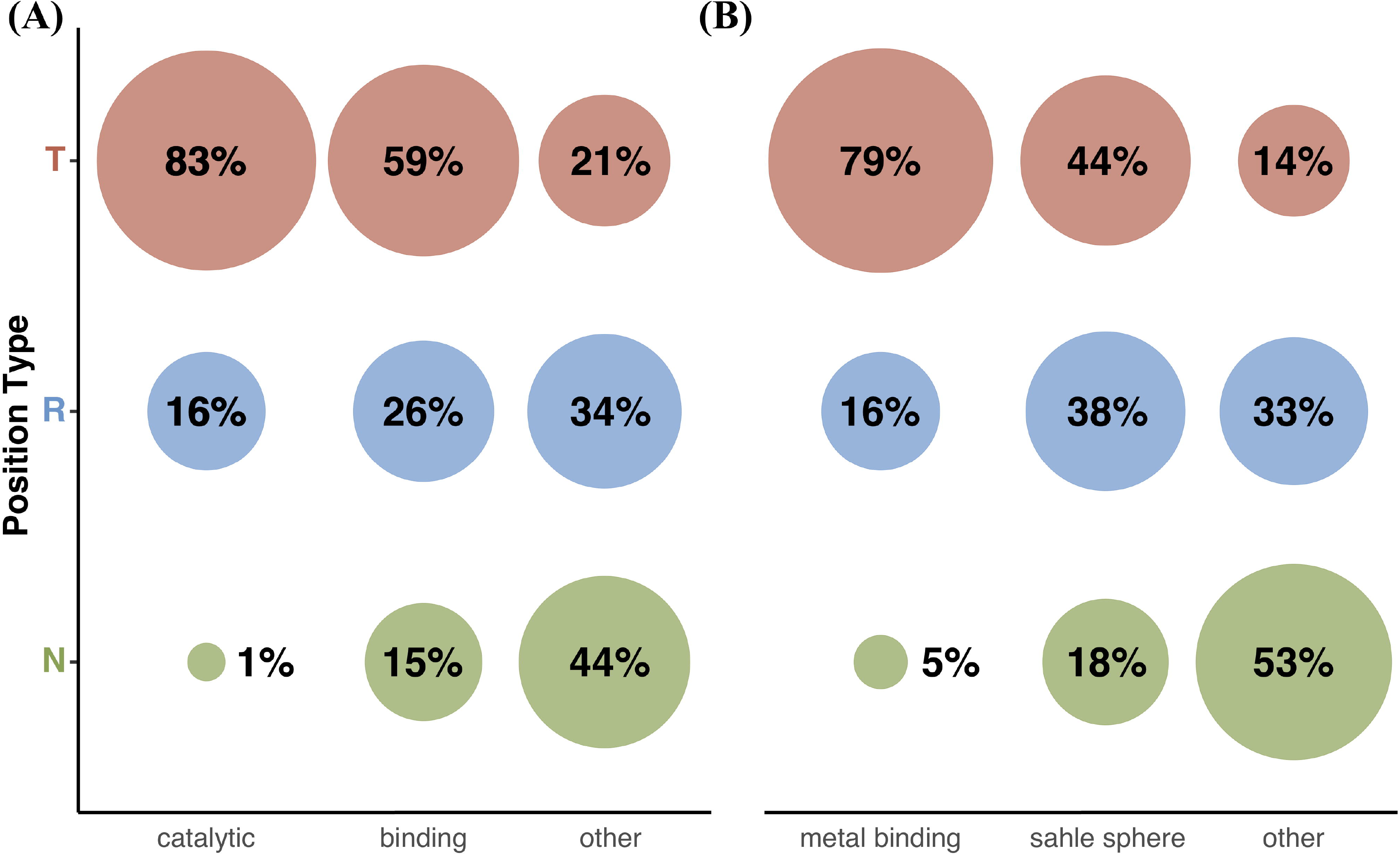
Distribution of position types across protein sites. (**A**) Enzymes, (**B**) Metal binding proteins. Colors are according to position type (green =*Neutral*, blue =*Rheostat*, red =*Toggle*). Percentages are rounded off and thus do not add up to 100%.

As expected, the majority of catalytic sites were *Toggles* and only 1% were *Neutral.* Binding sites were less frequently *Toggles* than catalytic sites, but much more frequently so than the *other residues* in the respective proteins, which were predominantly *Neutral*. Curiously, the fraction of *Rheostat* positions did not vary as drastically across the residues sets.

Notably the catalytic site primary actors – the charged amino acids (D, E, R, K, H; Supplementary Figure S4) (66) were unexpectedly low among the *Toggles* and *Rheostats* of the *other residues*; *i.e.* they were very important in functional sites, but not as relevant in other sites of the protein. This finding is particularly interesting in the light of the generic assumptions made about irreplaceability of charged residues. Outside the enzymatic functional sites, the more commonly structure-relevant large hydrophobic amino acids (C, W, Y, M, F) were most often *Toggles*, while the smaller (A, I, L, V) were drastically enriched in *Rheostats* (Supplementary Figure S4).

### Distribution of position types varies by metal-ligand binding proximity

We evaluated the composition of position types of residues located in the proximity of metal-containing ligands (*sahle* 3D-structure spheres, Methods) for Swiss-Prot proteins. As for functional sites above, we defined three sets of residues: those annotated in Swiss-Prot as metal binding, *sahle* sphere residues within 15A of the ligand center, and *other residues* (Figure 7B). Note that we excluded from consideration any residues annotated as metal binding and not located within a *sahle* sphere.

Metal binding residues showed a similar distribution of position types as catalytic sites (80% *Toggle*, 5% *Neutral*). Notably, *sahle* spheres were more enriched in *Rheostats* (38%) than were the binding sites described above (26%). However, the latter were more frequently Toggles (59%) than the former (44%). This result suggests that binding sites are critical features of function, while *sahle* spheres encompass residues relevant to functional flexibility. Moreover, outside of *sahle* spheres *Toggles* were the least abundant and more than half of the residues were *Neutral*, suggesting that most of the other residues are significantly less involved in protein function (including stability effects).

Preferred residues for metal binding are C, H, D, and E (67), which is also confirmed by our data (Supplementary Figure S5). Interestingly, *Toggles* were the dominant position type for all of these except glutamic acid (E), which were mostly *Neutrals* or *Rheostats*. One explanation for the observation, that the majority of variants only slightly impacts protein function, is the greater length of the glutamic acid side chain. This introduces a greater flexibility which allows for a larger range of possible substitutions than a more rigid structure.

### Position type profiles enable identification of disordered proteins

Based on MetaDisorder predictions (Methods) we labeled 6,309 Swiss-Prot proteins as disordered and 13,192 as ordered and compared the ratios of position types between these sets. The two classes of proteins were clearly separable by distribution of position types (Supplementary Figure S6).

Ordered proteins contained more than twice as many *Toggle*s as disordered proteins (19% vs. 8%), while disordered proteins were preferentially *Neutral* (68% vs. 46%). Of the 668 proteins, where *Neutrals* made up over 80% of all residues, 94% (650) were disordered. This result is, to a certain extent, expected due to frequent modulation of function, *i.e. Rheostatic* activity, achieved via structural changes; *e.g.* changes in residue solvent accessibility or secondary structure may, and often do, modulate functionality (68). However, this finding may also indicate that disordered proteins are poorly predicted by *funtrp*, as our method relies on structural features. Another hypothesis based on this observation may be that our definition of position types is not directly applicable to disordered proteins, where changes in functionality may be harder to objectively measure and evaluate.

### Experiments focus on high impact variants

We evaluated the relationship of position types with experimental annotations of variant effects extracted from the literature (PMD effect annotations as reported in (58)). Note that the number of PMD variants was the same across position types, *i.e.* 33% *Neutral*, 33% *Rheostats*, and 34% *Toggles*. On the other hand, position type ratios per protein are not at all similar on average, *i.e.* SwissProt had ~53% *Neutral* positions, ~33% *Rheostats*, and ~14% *Toggles*. This variant distribution difference suggests a strong preference in experimental studies towards evaluating the most likely non-neutral/severe variants and/or the most likely functionally or structurally important regions, in contrast to the unbiased DMS approach.

Based on PMD effect annotations, variants could be categorized into three main impact groups: *neutral*, *mild/moderate*, and *severe* (Supplementary Figure S7). In line with the above reasoning, there were more *severe* variants (43%) than *mild/moderate* variants (36%), and significantly more of either than of the *neutral* variants (20%). We further evaluated the PMD variant-affected position types. As expected, most of the *Toggle* positions (90%) had variants of at least some impact (severe or *mild/moderate*; 58% severe only). The fraction of *Rheostat* positions having non-neutral variants was slightly lower (80% any effect; 50% severe only). However, even as much as a third (35%) of the *Neutral* positions had severe variants (67% any impact). This high level of variant impacts across all evaluated protein positions underlines the exaggerated specific selection in experimental studies for expected-to-be-observed impact.

These observations of the bias in the reported variant impacts highlight the need for variant effect predictors to take into account or, at least, be mindful of, their effect-focused training/testing/development data.

### Position types can improve variant effect prediction

Changing the perspective to examine variant localization per position we observed that, as expected, roughly half (52%) of 3,254 *neutral* variants were in *Neutral* positions, but a full 18% of these affected *Toggles* (Supplementary Figure S7). Most (41%) of the *severe* (6,872) variants affected *Toggle* positions and were least abundant in *Neutrals* (25%). The variants in the *mild/moderate* group were nearly evenly distributed (33%, 32%, 35% *Neutrals*, *Rheostats*, *Toggles*, respectively) across all position types. Note that finding some *neutral* variants in *Toggle* positions and some *severe* variants in *Neutral* positions is not unexpected as per our position type definitions, which allow for some variety of effects. However, as *funtrp* has not been trained to recognize variant effects, the dominant trend of finding variants of expected impact in the right places, i.e. *neutral* variants in *Neutrals* and *severe* variants in *Toggles*, highlights our method’s ability to recognize functionally relevant protein positions.

The VarCards *Ensemble Prediction Ratio/Score* (Methods), which reflects the agreement of commonly used variant effect prediction tools, correlated with the severity of PMD impacts (Figure 8) across position types; *i.e.* there were more *non-neutral (mild/moderate and severe*) variants at higher scores than at lower scores, while the opposite was true for *neutral* variants. The recall of *neutral* variants was highest at *Neutral* positions (66% in *Neutral*, 44% in *Rheostats*, and 23% in *Toggles*; Figure 8 and Supplementary Table S6). In *Toggle* positions, *non-neutral* variants were more frequently incorrectly identified as *no-effect* than *neutral* variants (30% *no-effect* precision). These results suggest that *no-effect* predictions at *Toggle* positions are less reliable than at *Neutral* positions (F1_neutral_ =.48 and =.26, in *Neutrals* and *Toggles* respectively). As expected, prediction of *neutral* variants at *Rheostat* positions is less reliable than in *Neutrals*, but more reliable than in *Toggle* positions (F1_neutral_=.39).

**Figure 8.**
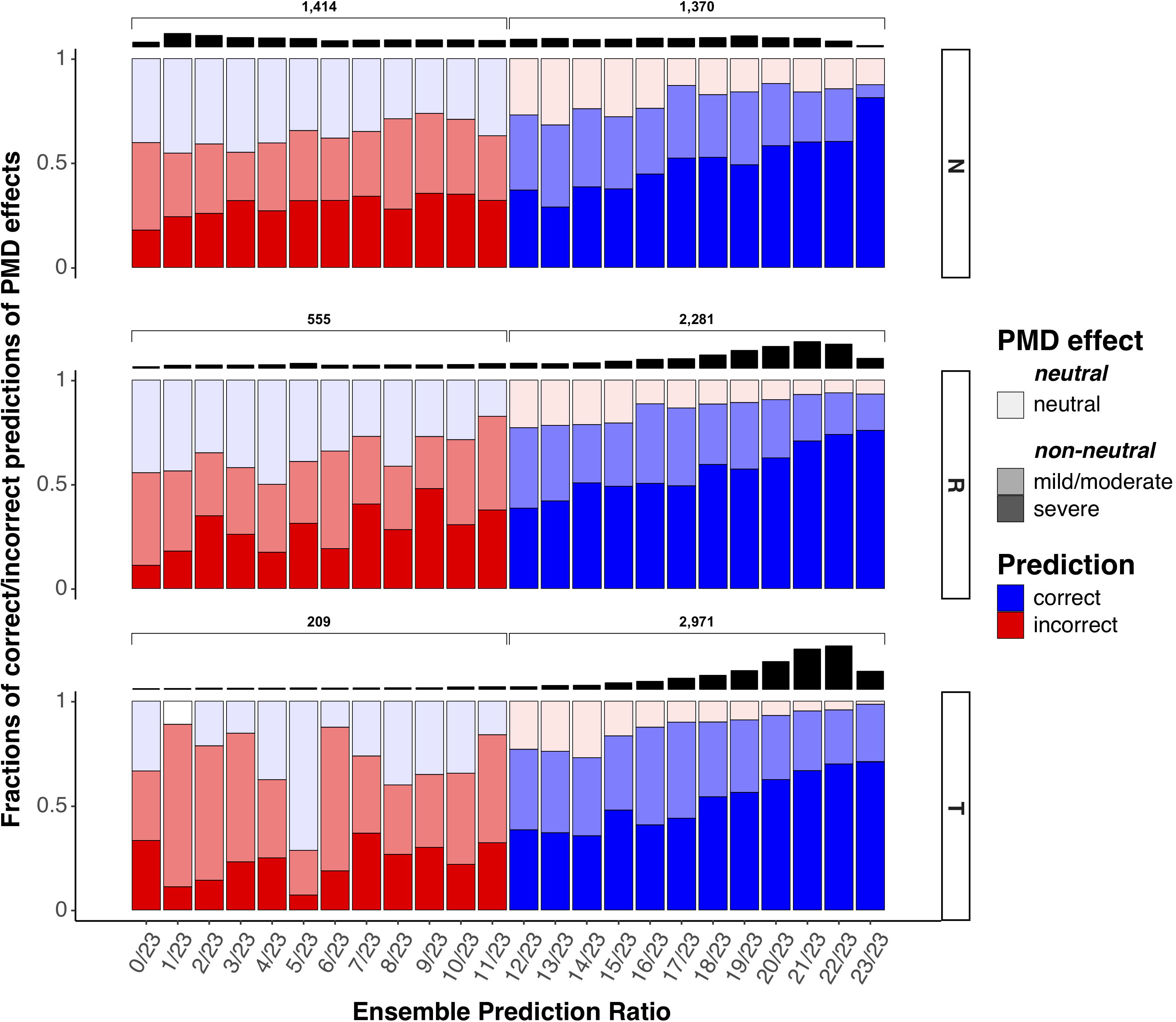
Fractions of correct and incorrect VarCards Ensemble predictions across PMD effect groups and *funtrp* position type. Each column reflects the fraction (y-axis) of correct (blue) and incorrect (red) Ensemble predictions (x-axis) per PMD effect group (light = neutral, medium = mild/moderate, dark = severe). The *Ensemble prediction ratio* signifies the fraction of tools predicting effect (*no-effect* prediction = 0/23, all methods predict *effect* = 23/23).

While predictions of *neutral* variants were worst at *Toggle* positions (77% incorrect), as a fraction of all *effect* predictions per position type, these errors were fewest (*effect* precision 93% in Toggles, 89% in *Rheostats* and 80% in *Neutrals*). *Non-neutral* recall was also highest in *Toggle* positions (95% in *Toggles*, 85% in *Rheostats*, and 56% in *Neutrals*), suggesting that *effect* predictions at *Toggle* positions are more reliable than at *Rheostat* or *Neutral* positions (F1_effect_ =.66, =.87, =.94 in Neutrals, Rheostats, and Toggles respectively)

To demonstrate the potential impact of position type knowledge on individual variant effect predictors we additionally evaluated the per position type performance for *effect* and *no-*effect predictions of SNAP, SIFT, PolyPhen-2 and Envision. As with VarCards scores, *Effect* predictions showed consistently higher performance at *Toggle* compared to *Neutral* positions (Figure 9, left panel in grey) while *no-effect* predictions showed consistently higher performance at *Neutral* compared to *Toggle* positions (Figure 9, right panel). Notably, this was the case for all three traditional variant effect predictions methods as well as the more recent DMS data trained approach. These findings unambiguously show that incorporating position types leads to much more reliable variant effect prediction.

**Figure 9.**
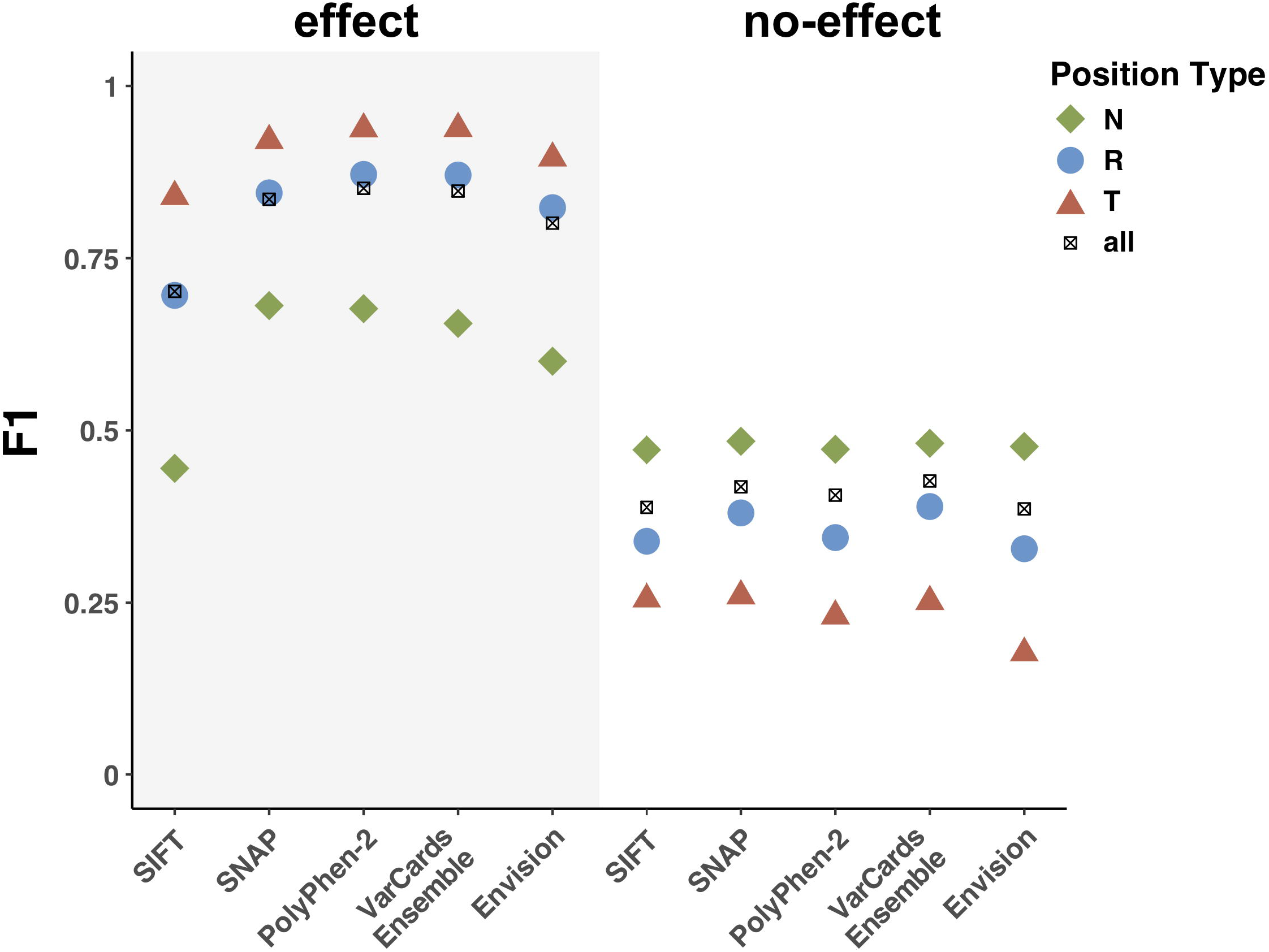
Performance of variant effect predictors significantly improves when considering relevant position type. Performance (F_1_ score) of *five* variant effect predictors for *effect* (grey, left panel) and *no-effect* (right panel) predictions of PMD variants, evaluated overall (black crosses =all) and per position type (green =*Neutral*, blue =*Rheostat*, red =*Toggle*).

To further highlight the relationship between *funtrp* position type predictions and annotated variant effects, we calculated the *no-effect vs. effect* ratios per type across a range of *funtrp* prediction scores (Figure 10). In line with the above results, we found that reliably predicted *Toggle* positions were more likely to have more *effect* variants (a lower ratio of *no-effect* to *effect* variants), while reliably predicted *Neutrals* had more *no-effect* variants (a higher ratio).

**Figure 10.**
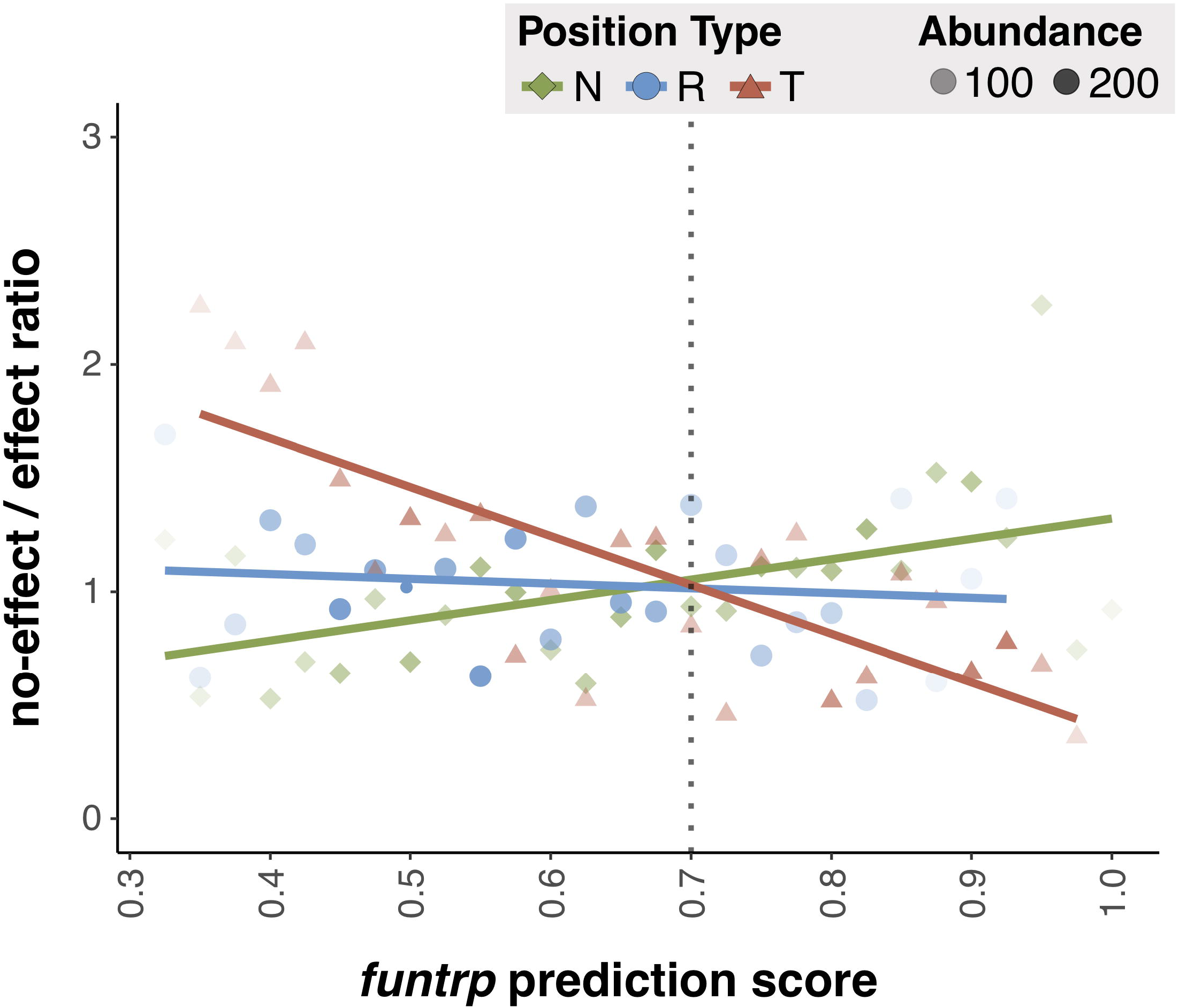
*funtrp* predictions correlate with PMD effect annotations. Ratio of *no-effect* vs. *effect* PMD variants (y-axis) per position type (green =*Neutral*, blue =*Rheostat*, red =*Toggle*) at respective *funtrp* prediction scores (x-axis). The abundance of position types at a certain score is represented by dot opacity. Trendlines are shown in the same color scheme.

Thus, we suggest that variant effect predictors could improve significantly if trained/developed separately for each *funtrp* position-type and/or accounting for the reliability of position type prediction.

#### An aside

We expect that prediction could be most improved for *Rheostats*, where increased resolution is likely once the obvious *Toggle* and *Neutral* variants are no longer the main focus. We note that *Rheostat* positions are the most likely proverbial fireplaces for burning the fuel for evolution (58), *i.e.* introducing a multitude of tiny changes to optimize a functionality best fit to the particular environment. Tracing the conversion of *Rheostats* into *Neutrals* or *Toggles* across homologs can therefore highlight the evolutionary paths taken or currently in place for any given molecular functionality. Thus, our new definition of position types will likely contribute to the understanding of biophysics of protein folding and related epistatic mutation effects, highlight prime candidate residues for directed evolutionary pathways, and help shine light on pathogenicity mechanisms.

## AVAILABILITY

The Java based implementation of Random Forest Classification is part of the WEKA library (60) and can be found at https://sourceforge.net/projects/weka.

R is is a free software environment for statistical computing and graphics and can be downloaded at https://www.r-project.org/.

The PredictProtein docker image used in this work can be found at https://doi.org/10.5281/zenodo.3018245. The latest release is available via Docker Hub (bromberglab/predictprotein). The source code is accessible at https://bitbucket.org/bromberglab/predictprotein.

The *funtrp* prediction pipeline docker image used in this work can be found at https://doi.org/10.5281/zenodo.3020352. The latest release is available via Docker Hub (bromberglab/funtrp). The source code is accessible at https://bitbucket.org/bromberglab/funtrp. *funtrp* training data is also available and can be found at https://doi.org/10.5281/zenodo.3066344.

*funtrp* is also available as webservice at https://services.bromberglab.org/funtrp.

## SUPPLEMENTARY DATA

Supplementary Data are available at bioRxiv online.

## ACKNOWLEDGEMENT

We would like to thank Dr. Liskin Swint-Kruse (University of Kansas) for all help and comments that made this work possible. We are also grateful to Dr. Predrag Radivojac (Northeastern), Dr. Jay A. Tischfield, Dr. Gary Heiman, Dr. Chengsheng Zhu, Yannick Mahlich, Yanran Wang, and Zishuo Zeng (all Rutgers) for all discussions and to Dr. Sonakshi Bhattacharjee (Columbia) for discussions and help with the manuscript. We would also like to express gratitude to all people who deposit their data into publicly available databases and to those who maintain them.

