## Supplementary Figure for "fuNTRp: Identifying protein positions for variation driven functional tuning"

**Table S1.** Training dataset composition

|  | *Toggle* | *Neutral* | *Rheostat* | SNP-Possible | unknown | filtered | detailed |
| --- | --- | --- | --- | --- | --- | --- | --- |
| raw | - | - | - | - | 822 | - |  |
| X-residues |  |  |  |  | 820 | 2 | undetermined amino acid X |
| SNP-possible | - | - | - | 591 | 229 | - |  |
| k-means labeling | 66 | 152 | - | 372 | 229 | - | k-means clustering exp. scores |
| manual refinement | 60 | 147 | - | 372 | 229 | 12 | removed: T (6), N (6) |
| ntModel labels | 94 | 181 | 151 | 61 | 113 | - | removed: T (74), N (109), R (48) |
| manual refinement | 20 | 72 | 104 | - | - | 231 |  |
| *funtrpTraining* | 80 | 219 | 104 | 61 | 113 | 245 | Final *funtrpTraining* = 403 |

Detailed overview of the of training dataset composition for training of both random forest-based models in the funtrp pipeline. Show are the total numbers of instances remaining after the cluster labeling, filtering and prediction steps.

**Table S2.** Set of sequence-based features used to train funtrp random forest-based models

| id | Feature | Source | Description | Parameters |
| --- | --- | --- | --- | --- |
| 1 | Solvent Accessibility | PROF (*) | predicted solvent accessibility (PACC) | PredictProtein defaults |
| 2 | Secondary Structure | PROF (*) | predicted helix (pH), strand (pE) or loop (pL) | PredictProtein defaults |
| 3 | Residue Flexibility | PROFbval (*) | predicted residue flexibility (PROFbval) | PredictProtein defaults |
| 4 | Protein Disorder | MD (*) | predicted protein disorder (MDraw) | PredictProtein defaults |
| 5 | Amino Acid | - | amino acids encoded as a vector of length 20 | NA |
| 6 | Residue Size | - | basic amino acid property (small or large) | NA |
| 7 | Residue Charge | - | basic amino acid property (uncharged / + / -) | NA |
| 8 | SNP possible | - | number of possible nsSNPs (all codons) | NA |
| 9 | Conservation | ConSurf (*) | predicted conservation | PredictProtein defaults |
| 10 | MSA Ratio | - | Total fraction of residue amino acid at MSA column | NA |

(*) tools are applied via the PredictProtein pipeline (Yachdav et al., 2014). Features were ranked by importance towards fuNTRp position type labels in Swiss-Prot using ReliefF; weights were rounded off (Kononenko, RobnikSikonja, & Pompe, 1996). If applicable, parameters used in feature computation are specified.

| Identifier | Source | Proteins <> *fuNTRp* | w/ E.C. annotation | w/o E.C. annotation |
| --- | --- | --- | --- | --- |
| Swiss-Prot | UniProtKB/SwissProt | 20,410 <> 19,501 | 4,273 <> 4,241 | 16,137 <> 15,260 |
| TrEMBL | UniProtKB/TrEBML |  | 9,668 <> 9,554 | 144,277 <> 5,254 |
| EXPV | UniProtKB/SwissProt | 1,250 <> 1,239 | 1,250 <> 1,239 | x |
| PMD | PMD & (Bromberg, Kahn, & Rost, 2013) | 1.224 <> 1,220 | x | x |

**Table S3.** Protein subsets for model training

Extracted datasets used in analysis. EXPV is a subset of experimentally verified enzymes in Swiss-Prot (Mahlich, Steinegger, Rost, & Bromberg, 2018). Literature based annotations of effect (PMD database) were taken from (Bromberg et al., 2013).

**Table S4.** Confusion matrices of position type predictions for (A) ntModel und (B) funtrpModel

**(B)**

**(A)**

| *Neutral* | *Toggle* | *Rheostat* | Observed ↓ |
| --- | --- | --- | --- |
| 199 | 4 | 16 | *Neutral* |
| 2 | 64 | 14 | *Toggle* |
| 19 | 5 | 80 | *Rheostat* |

| *Neutral* | *Toggle* | Observed ↓ |
| --- | --- | --- |
| 140 | 7 | *Neutral* |
| 9 | 51 | *Toggle* |

Predictions for both models are based on LOO-CV results.

**Table S5.** Performance of predicting position types for a Random Forest (RF) based classifier model using evolutionary conservation alone

| Position Type | Sensitivity | Specificity | PPV | NPV | Precision | Recall | F1 | Prevalence | Detection Rate | Detection Prevalence | Balanced Accuracy |
| --- | --- | --- | --- | --- | --- | --- | --- | --- | --- | --- | --- |
| *Neutral* | 0.66 | 0.87 | 0.75 | 0.81 | 0.75 | 0.66 | 0.70 | 0.38 | 0.25 | 0.33 | 0.76 |
| *Rheostat* | 0.46 | 0.70 | 0.29 | 0.83 | 0.29 | 0.46 | 0.35 | 0.21 | 0.10 | 0.33 | 0.58 |
| *Toggle* | 0.66 | 0.89 | 0.81 | 0.79 | 0.81 | 0.66 | 0.72 | 0.41 | 0.27 | 0.33 | 0.77 |

Shown are the averaged performances per class over 100 resample runs. For each run, 3000 residue positions from Swiss-Prot were resampled randomly (without replacement), selecting 1000 instances of each position type respectively. The same was repeated for the test set and a total of 300 residue positions. Position type labels were based on *funtrp* predictions. PPV = positive predictive value; NPV = negative predictive value.

**Table S6.** Performance of VarCards Ensemble prediction for PMD effect annotation dataset

|  |  | **neutral** | **non-neutral** | |  |  |
| --- | --- | --- | --- | --- | --- | --- |
| **Position Type** | **VarCards prediction** | (neutral) | (mild/moderate | severe) | **total per effect** | **total per position type** |
| ***Neutral*** | **effect** | 273 | 445 | 652 | *1,370* | **2,784** |
|  | **no-effect** | 535 | 465 | 414 | *1,414* |  |
| ***Rheostat*** | **effect** | 245 | 624 | 1,412 | *2,281* | **2,836** |
|  | **no-effect** | 191 | 198 | 166 | *555* |  |
| ***Toggle*** | **effect** | 217 | 941 | 1,813 | *2,971* | **3,180** |
|  | **no-effect** | 63 | 94 | 52 | *209* |  |
|  | **total *Neutral*** | *808* | *910* | *1,066* |  |  |
|  | **total *Rheostat*** | *436* | *822* | *1,578* |  |  |
|  | **total *Toggel*** | *280* | *1,035* | *1,865* |  |  |
| **total per PMD effect** | | **1,524** | **2,767** | **4,509** |  |  |

**SPG1**

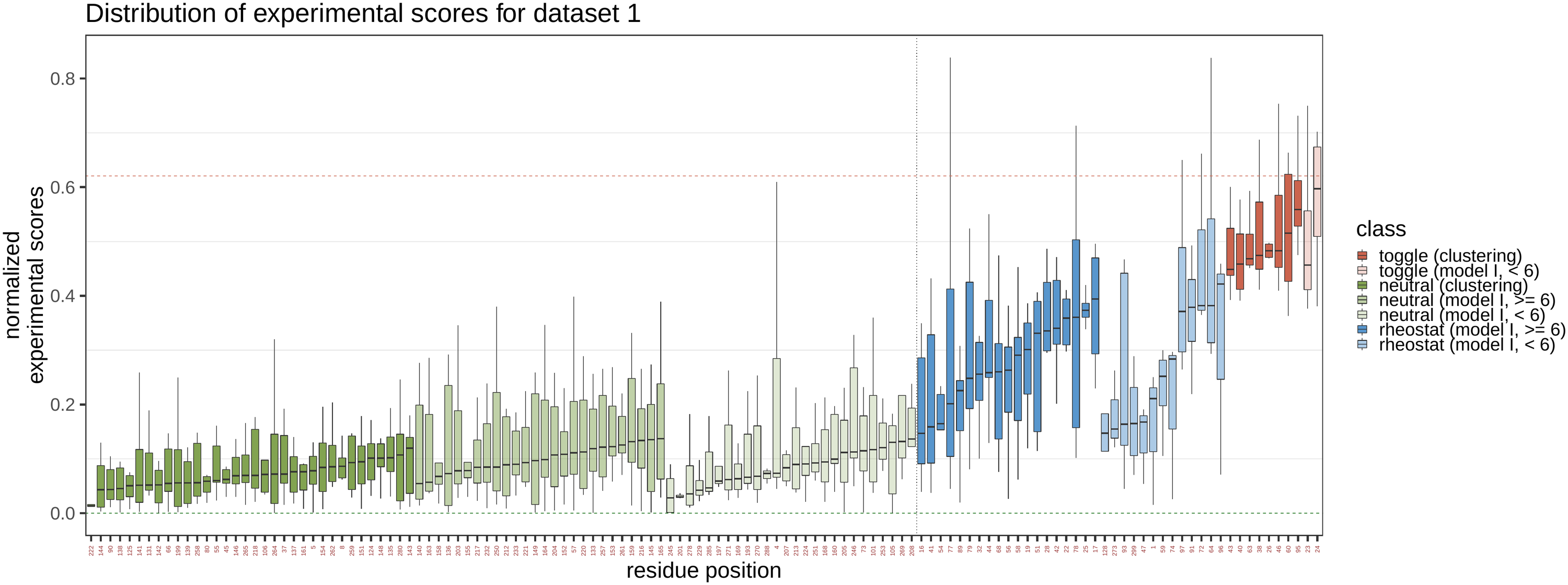

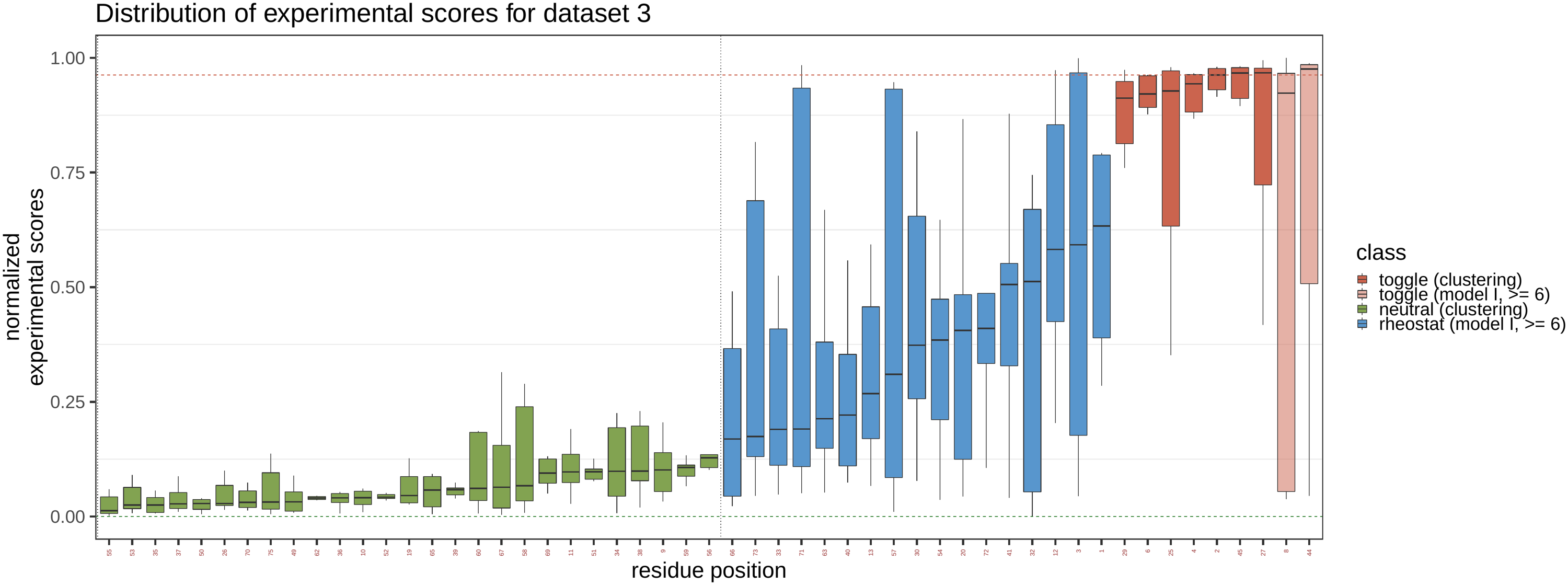

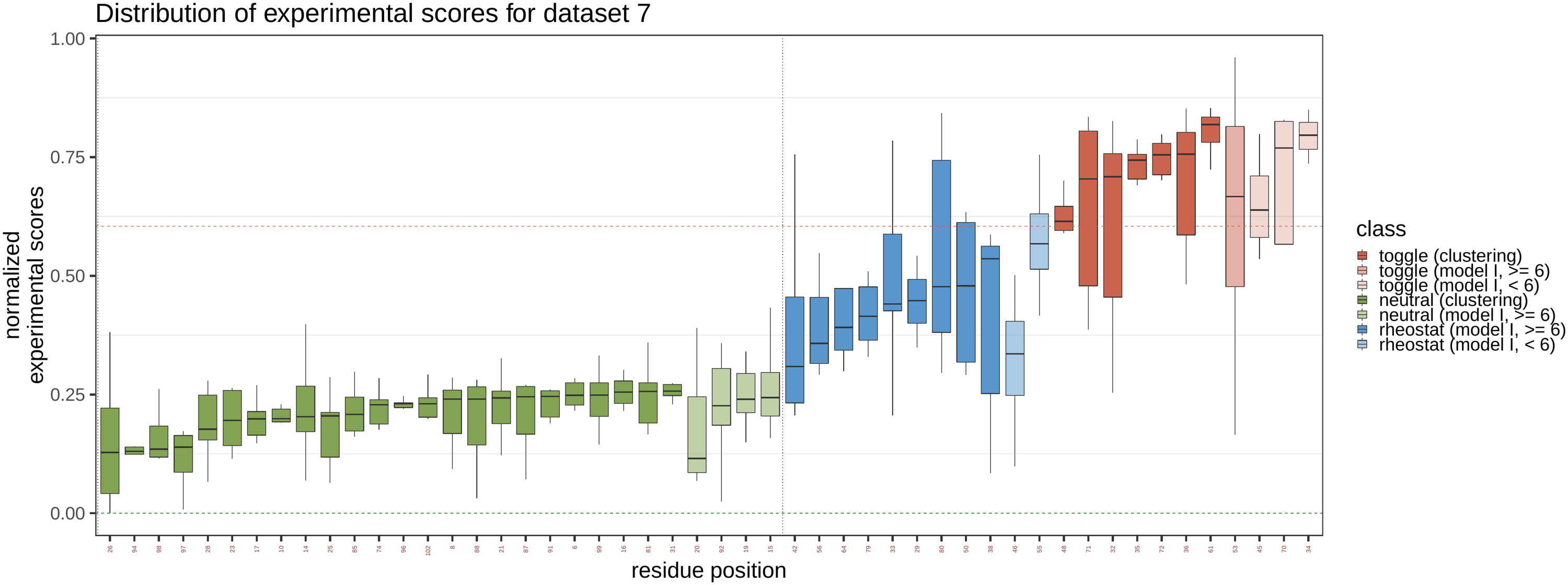

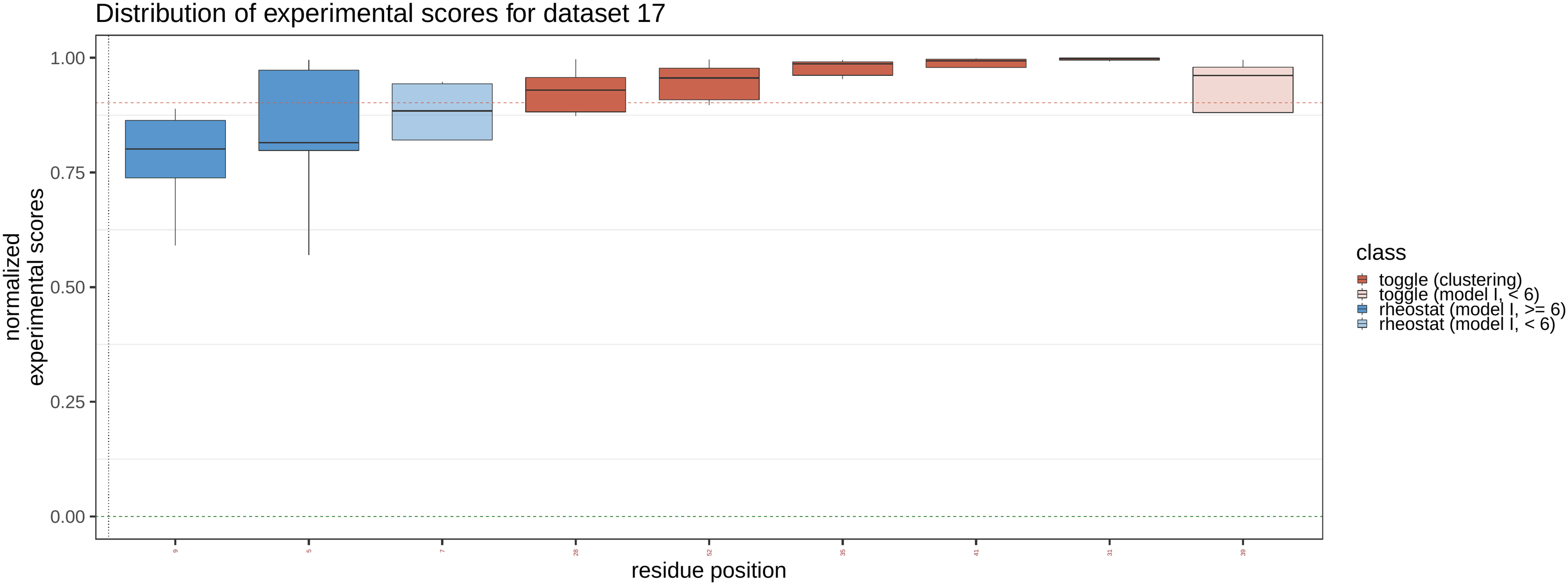

**Figure S1. Distribution of experimental (DMS) variant effect scores for training datasets.** Measured experimental scores extracted from DMS datasets were normalized to [0,1]. Residue positions on the x-Axis are grouped by (i) position types, (ii) way of labeling and (iii) within these groupings ordered based on increasing distribution medians. The labeling types are: clustering, predicted with more than six experimental scores available and predicted with less than six experimental scores available.

**UBE4B**

**PAB1**

**BRCA1**

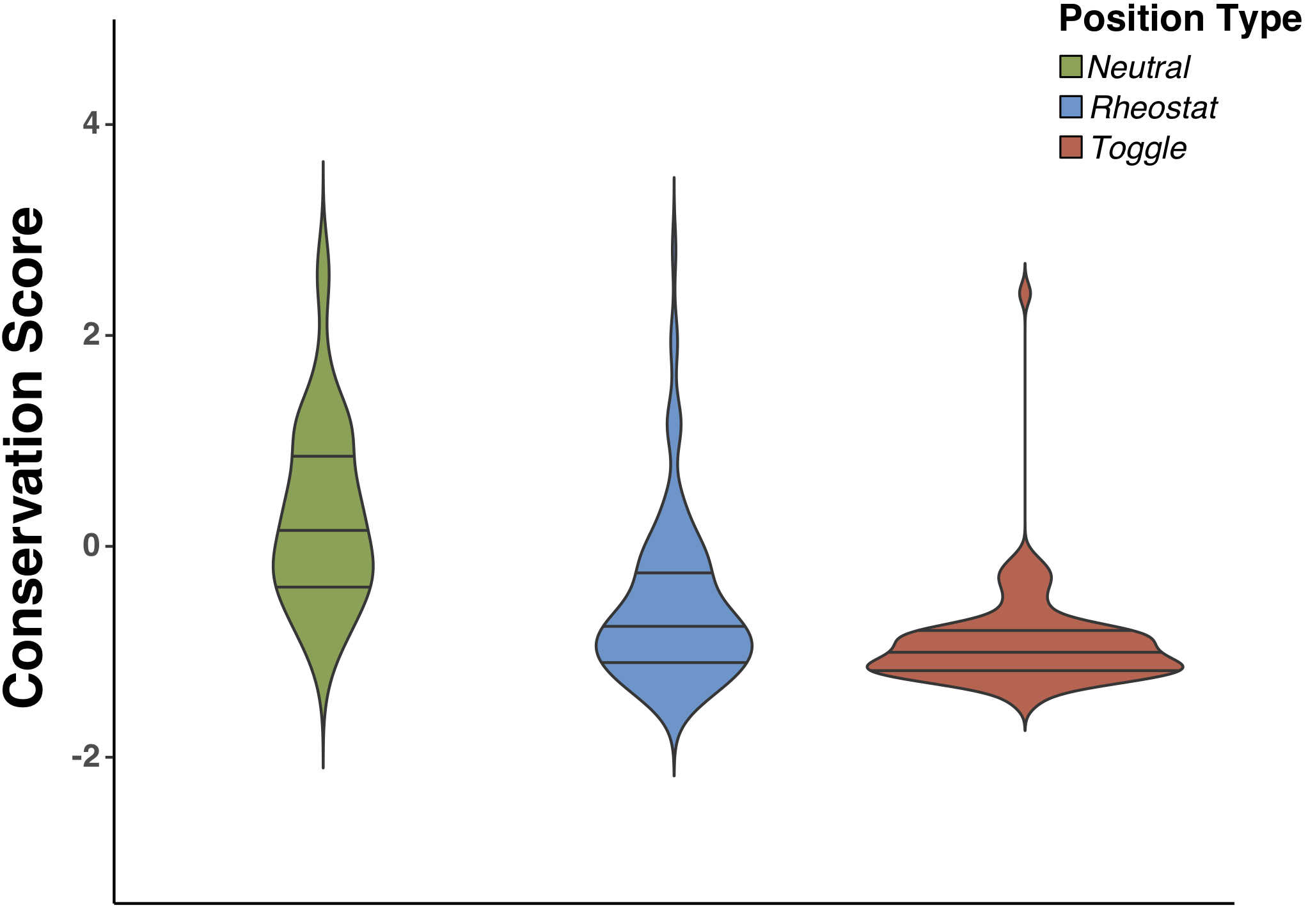

**Figure S2. Distribution of ConSurf conservation scores for fuNTRp training dataset.** Density distributions of evolutionary conservation (ConSurf) compared between position types for the *funtro* model training dataset. ConSurf predictions scores are by default normalized such as 0 depicts the average score over the entire protein and standard devia-tion is |1|). Colors are according to position type (green =*Neutral*, blue =*Rheostat*, red =*Toggle*).

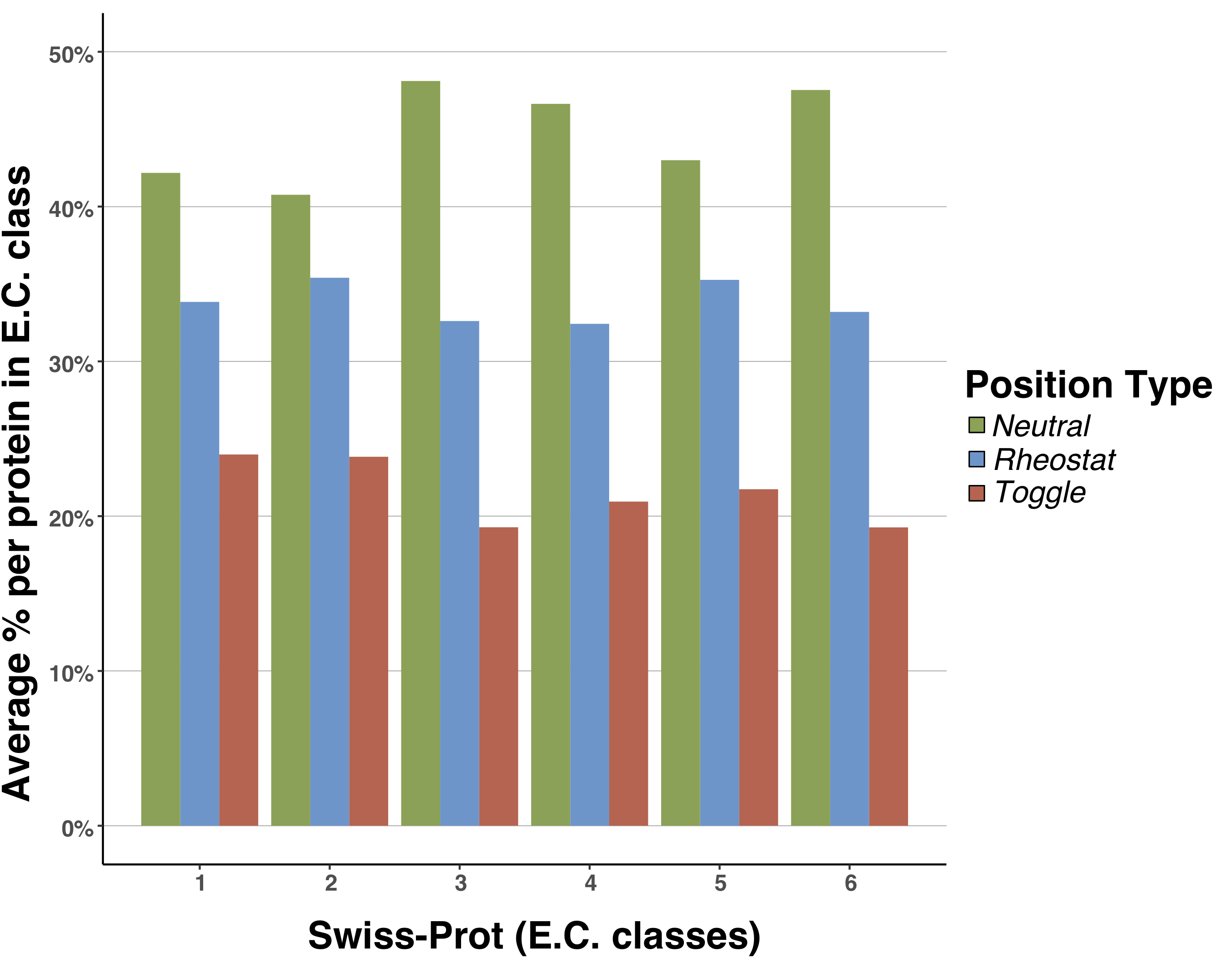

**Figure S3. Average fraction of position types on per-protein basis for main E.C. classes in the entire Swiss-Prot dataset.** Colors are according to position type (green =*Neutral*, blue =*Rheostat*, red =*Toggle*). Mean fractions of position types differ significantly among enzyme classes based on the standard error of the mean: 1 (N= 6.0E-04, R=6.4E-04, T=5.8E-04), 2 (N=3.7E-04, R=4.1E-04, T=2.4E-04), 3 (N=5.2E-04, R=4.2E-04, T=3.6E-04), 4 (N=9.2E-04, R=1.0E-03, T=7.6E-04), 5 (N=1.5E-03, R=1.1E-03, T=1.2E-03), 6 (N=8.9E-04, R=9.5E-04, T=9.1E-04).

**Figure S4. Fractions of position types per amino acid compared by site characteristic.** Comparison of fractions at catalytic sites and binding sites against the remaining residues of the respective Swiss-Prot enzymes. Colors are according to position type (green =Neutral, blue =Rheostat, red =Toggle).

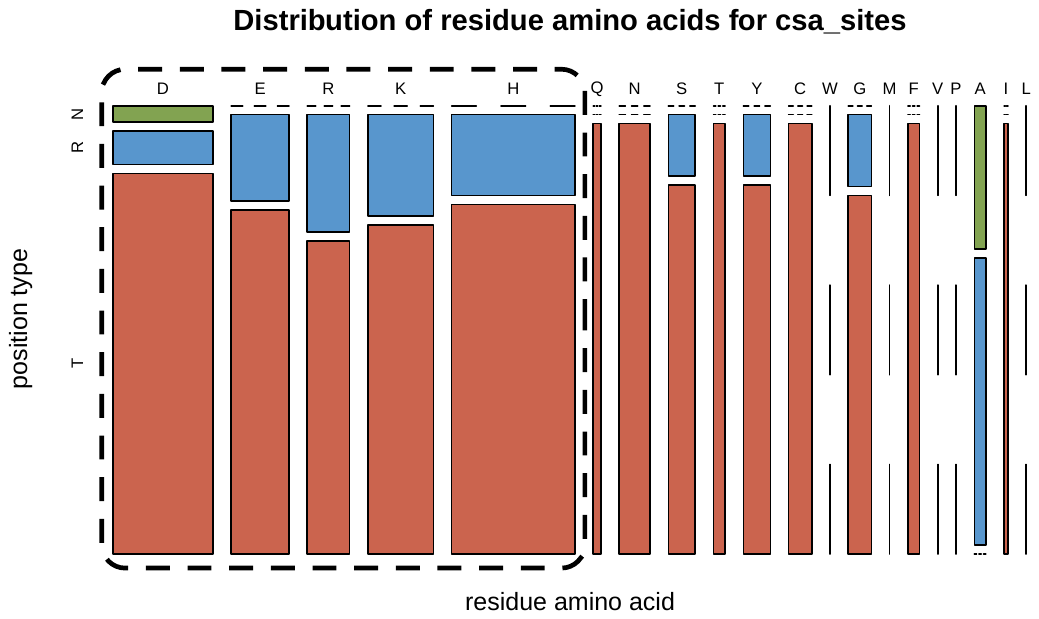

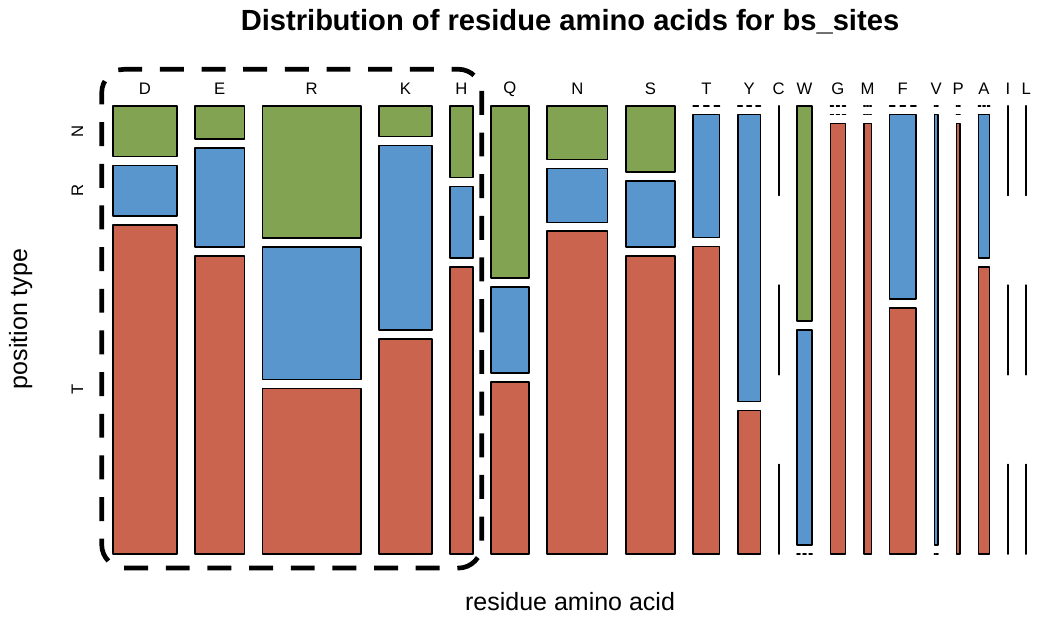

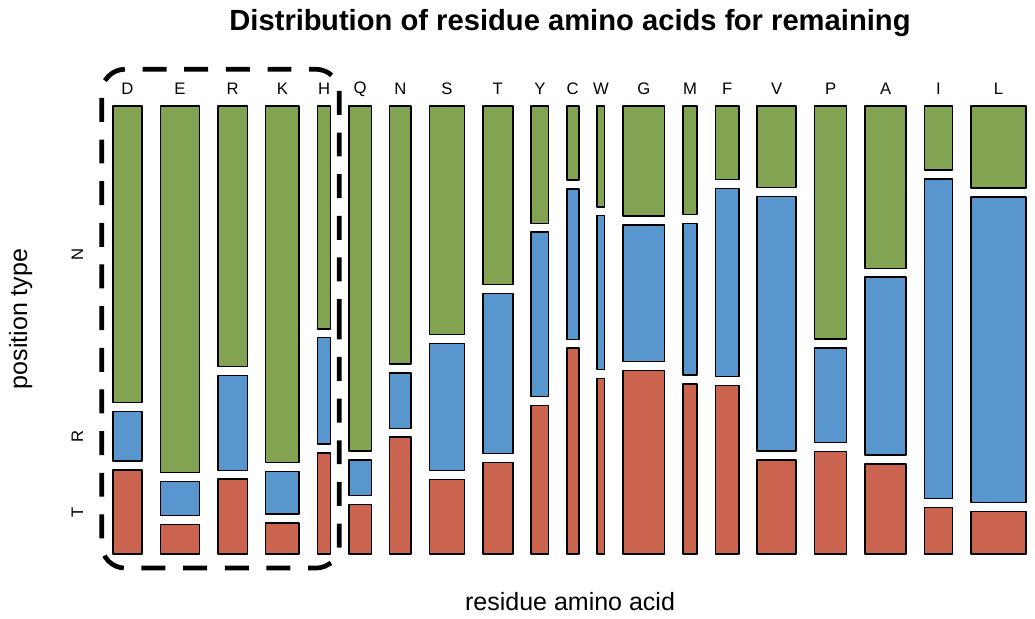

Polar, charged

Polar, charged

Polar, charged

Polar, hydrophilic

Non-polar, hydrophobic

Polar, hydrophilic

Non-polar, hydrophobic

Polar, hydrophilic

Non-polar, hydrophobic

Amino Acid and Group

Position Type at

catalytic sites

binding sites

remaining sites

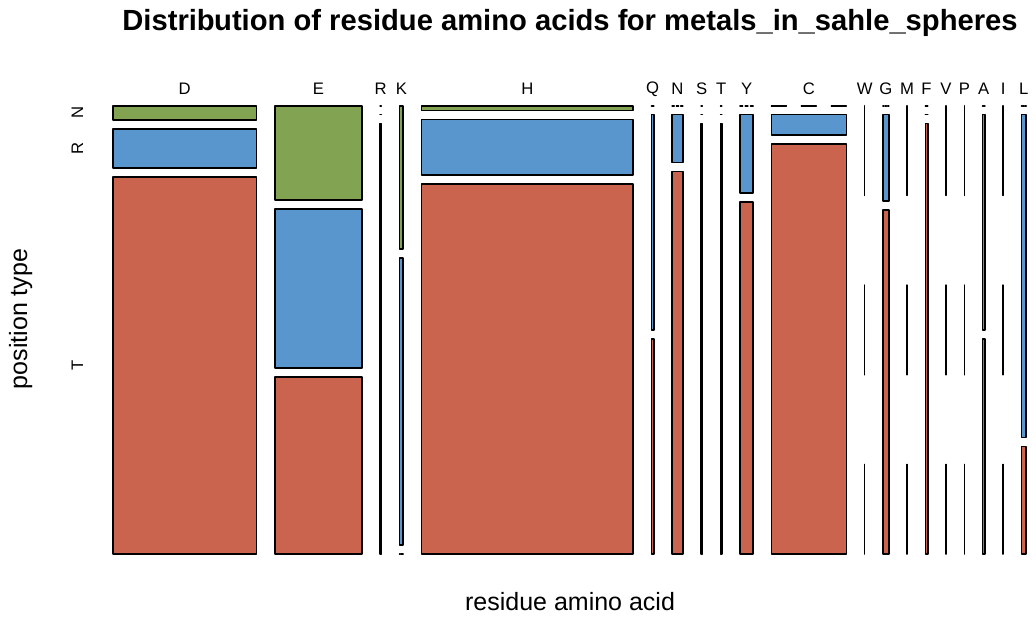

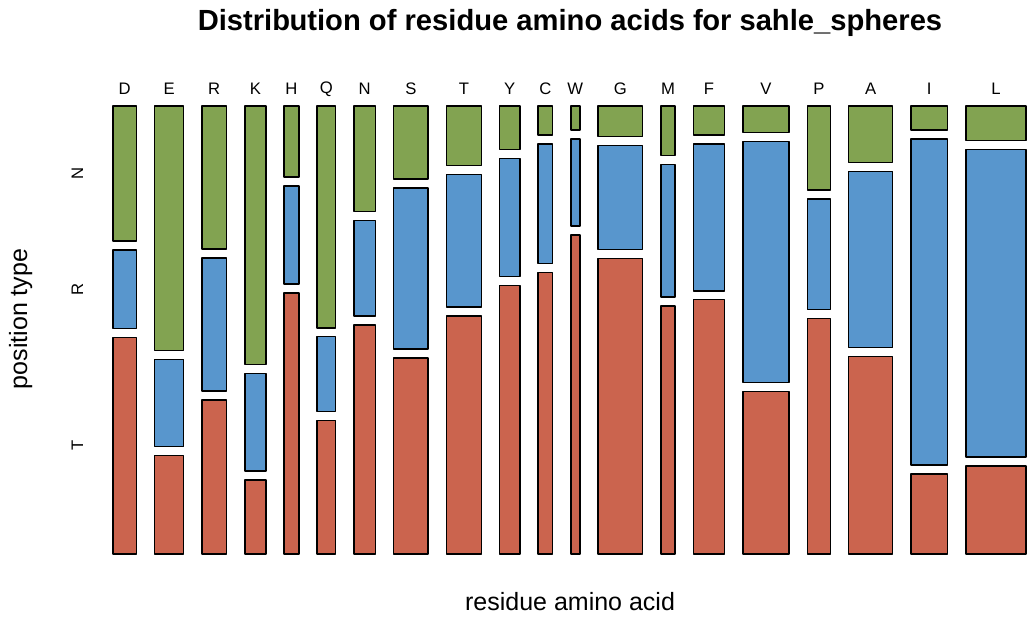

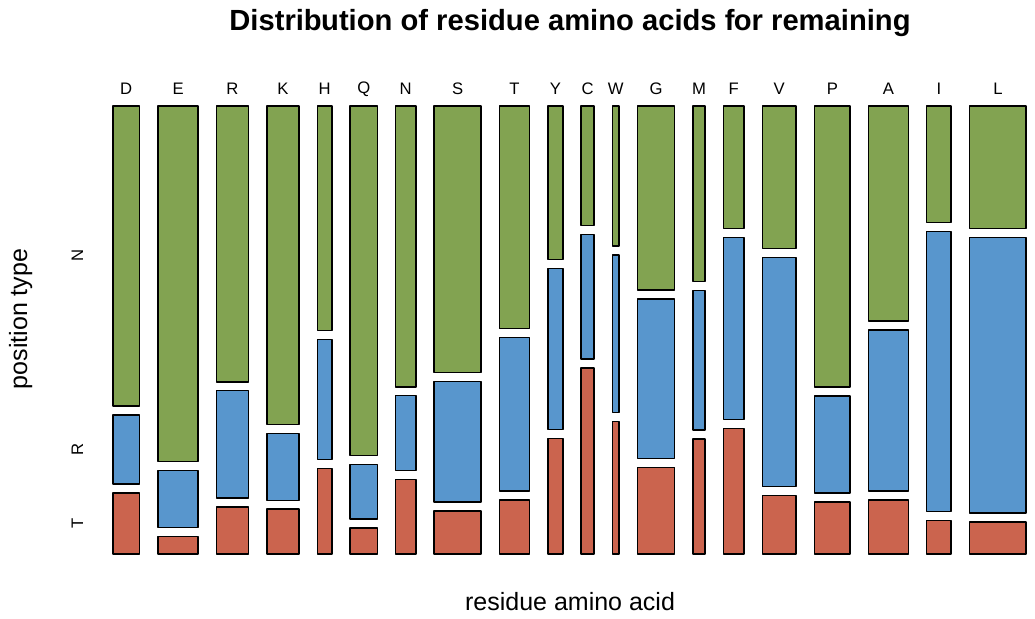

Position Type

Polar, charged

Polar, charged

Polar, charged

Polar, hydrophilic

Non-polar, hydrophobic

Polar, hydrophilic

Non-polar, hydrophobic

Polar, hydrophilic

Non-polar, hydrophobic

Amino Acid and Group

**Figure S5. Fractions of position types per amino acid for metal binding sites and spheres.** Comparison of SaHLe spheres and residues annotated as metal binding sites within spheres vs remaining residues of the respective Swiss-Prot enzymes. Colors are according to position type (green =Neutral, blue =Rheostat, red =Toggle).

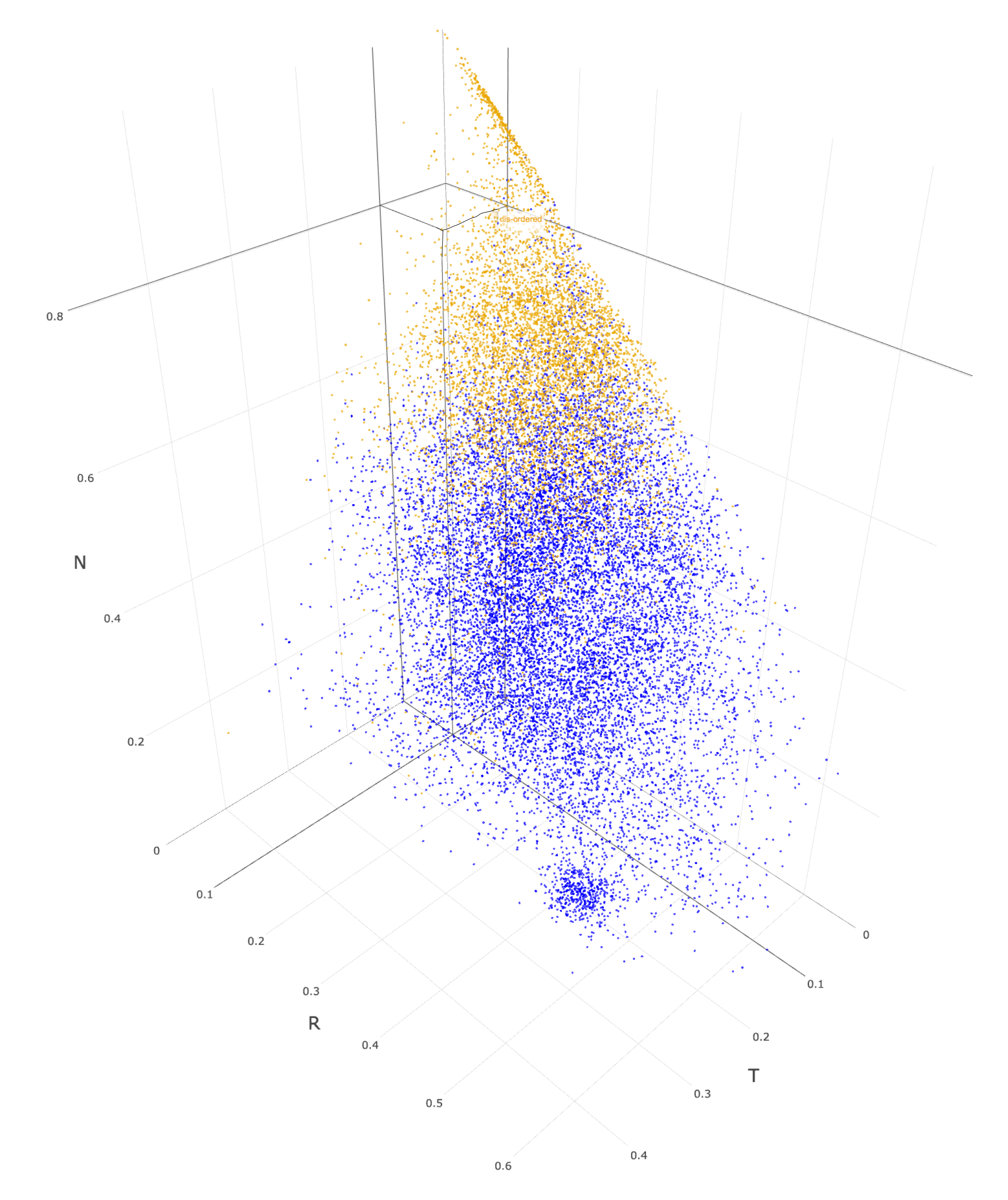

**Figure S6. funtrp prediction scores for disordered Proteins compared within position types.** Proteins in Swiss-Prot were labeled as either ordered or disordered based on MetaDisorder predictions (Methods). Residues located in disordered proteins are highlighted in yellow, those found in ordered proteins are shown in blue.

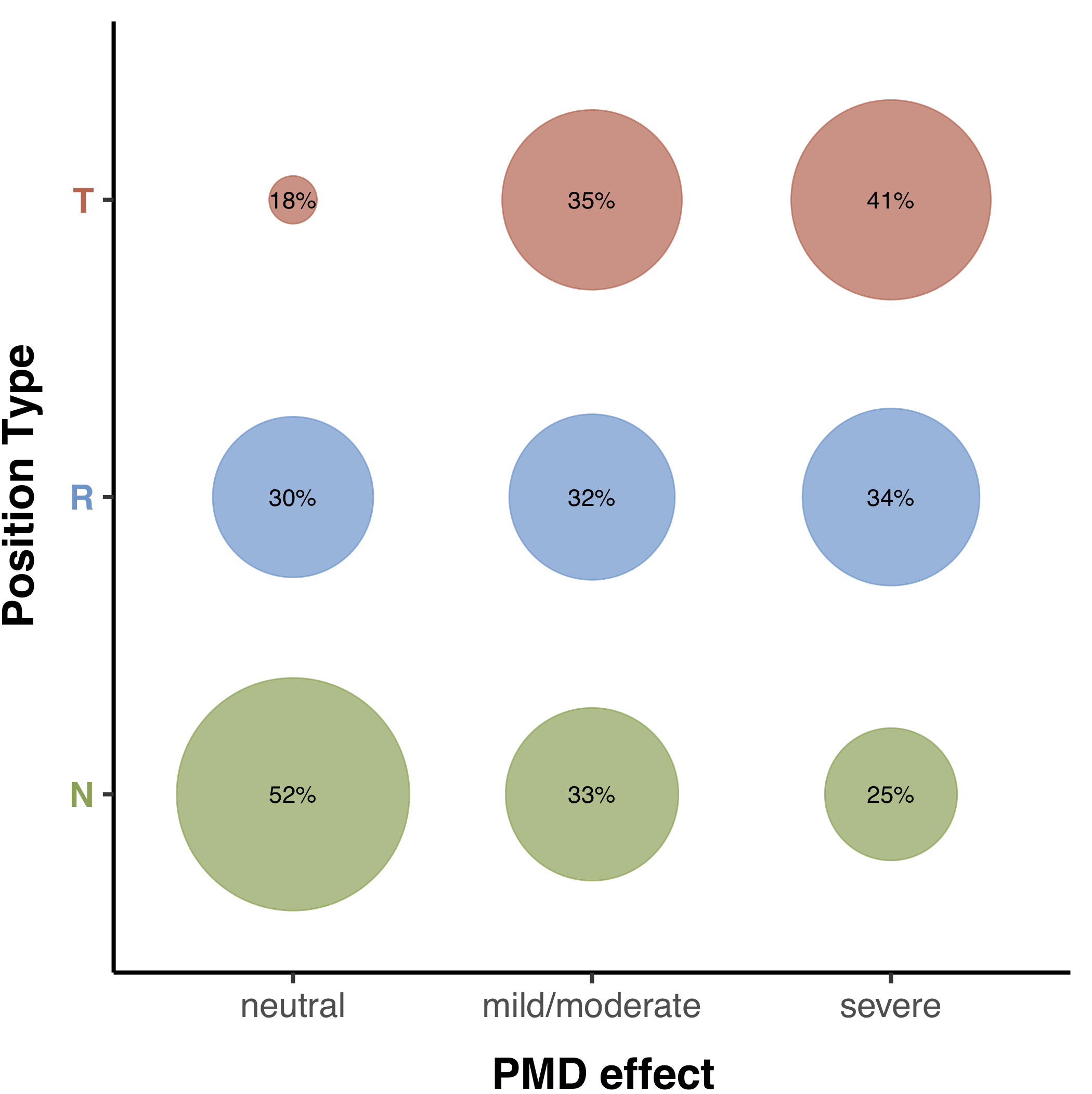

**Figure S7. Distribution of position types for PMD effect annotations.** PMD *mild* and *moderate* effects annotations were grouped into *mild/moderate*. Percentages are rounded; colors are according to position type (green =*Neutral*, blue =*Rheostat*, red =*Toggle*).
