## Supplementary Table S1 for "fuNTRp: Identifying protein positions for variation driven functional tuning"

**Table 1.** Deep mutational scanning datasets used in *funtrpModel* training and testing

| **Gene** | **Sub-region** | **Organism** | **Variants** | **Measured Activity** | **Set** |
| --- | --- | --- | --- | --- | --- |
| BRCA1 | RING domain | H. sapiens | 3,080 | Ubiquitin ligase activity | train |
| PAB1 | RRM domain | S. cerevisiae | 1,188 | mRNA binding specificity | train |
| UBE4B | U-box domain | H. sapiens | 926 | Ubiquitin ligase activity | train |
| TEM-1 | - | E. coli | 5,469 | Ampicillin resistance | train |
| SPG1 | GB1 | Strepto. sp | 467 | Binding affinity to IgG | train |
| PTEN | - | H. sapiens | 3,880 | Protein stability | test |
| TPMT | - | H. sapiens | 3,756 | Protein stability | test |
| HSP90 | ATPase domain | S. cerevisiae | 4,231 | Yeast growth | test |
