## Supplementary Table S2 for "fuNTRp: Identifying protein positions for variation driven functional tuning"

**Table 2.** Set of sequence-based features used by prediction model

| **id** | **Feature** | **Source** | **ReliefF**** | **Rank** |
| --- | --- | --- | --- | --- |
| 1 | Solvent Accessibility | PROF (*) | 0.18 | 3 |
| 2 | Secondary Structure | PROF (*) | 0.12 | 6 |
| 3 | Residue Flexibility | PROFbval (*) | 0.15 | 4 |
| 4 | Protein Disorder | MD (*) | 0.22 | 2 |
| 5 | Amino Acid | - | 5e-5 | 8 |
| 6 | Residue Size | - | 0 | 10 |
| 7 | Residue Charge | - | 1e-7 | 9 |
| 8 | SNP possible | - | 7e-4 | 7 |
| 9 | Conservation | ConSurf (*) | 0.34 | 1 |
| 10 | MSA Ratio | - | 0.14 | 5 |

(*) tools in the PredictProtein pipeline (Yachdav et al., 2014). (**) Features ranked by importance using ReliefF (Kononenko, RobnikSikonja, & Pompe, 1996). Secondary structure scores were reported per position for helix, sheet, and loops (pH, pE, and pL). Feature descriptions and default parameters are detailed in Supplementary Table S2.
